# ResLit: A Large-Scale Automated Literature Mining Database for Antimicrobial Resistance

**DOI:** 10.64898/2026.08.14.744991

**Authors:** A. Skoulakis, H. Xiao, K.A. Provatas, A. Galaras, G. A. Pavlopoulos, I. Georgakopoulos-Soares

## Abstract

Antimicrobial resistance generates a vast, rapidly growing literature, yet no resource offers a comprehensive, evidence-linked repository of AMR findings at scale. We present ResLit, an automated pipeline and public database that mines the AMR literature for resistance genes, mutations, organisms, and mechanisms. From 2 million candidate PubMed records, BioMistral-7B screened abstracts to 356,000 relevant papers; multi-tier retrieval yielded 117,000 full texts, from which Qwen3-30B performed two-step extraction. ResLit contains 3,120 genes and 13,593 mutations, cross-linked to CARD, ResFinder, and NCBI Reference Gene Catalog across four evidence tiers. It further supports community-driven curation of automated outputs and reference databases. Freely available at www.reslit.info.

## Background

Antimicrobial resistance (AMR) is a major and growing threat to global health, with bacterial AMR directly responsible for an estimated 1.14 million deaths in 2021. AMR is projected to cause 1.91 million deaths annually by 2050 **(GBD 2021 Antimicrobial Resistance Collaborators, 2024)**. The molecular mechanisms of resistance, including acquired genes, chromosomal mutations, efflux systems, target modifications, and enzymatic inactivation, have been extensively studied in the primary scientific literature, with a huge corpus of new studies published annually **(Blair et al., 2015; McArthur et al., 2013)**. The volume of this literature presents a significant challenge: researchers must integrate findings across organisms, antibiotics, geographic contexts, and experimental settings **(Edalatmand and McArthur 2023),** while curators of reference databases such as the Comprehensive Antibiotic Resistance Database (CARD) **(McArthur et al. 2013)**, ResFinder **(Bortolaia et al. 2020)**, and the NCBI Reference Gene Catalog (NCBI RGC) must identify the published papers and validate their results to curate entries.

Several curated AMR databases have substantially contributed to this field. CARD offers an ontology-driven, literature-linked resource of resistance determinants; ResFinder supports genotype-to-phenotype resistance prediction for a smaller, species-focused panel; and the NCBI Reference Gene Catalog provides a broad AMR reference dataset. Despite their utility, these databases rely on expert curation, a time-intensive process that inherently lags behind primary literature, particularly given the exponential growth in AMR publications. In addition, their coverage is still restricted to a relatively small subset of species, highlighting the need for broader reference frameworks to support AMR analysis across thousands of bacterial genomes **(Skoulakis et al., 2025)**.

Natural language processing (NLP) and large language models (LLMs) can accelerate literature scanning and synthesis. Recent advances in biomedical language models, including domain-specific models such as BioMistral **(Labrak et al. 2024)** and PubMedBERT **(Gu et al. 2020)**, as well as general-purpose instruction-following models such as Qwen3 **(Yang et al. 2025)**, have demonstrated strong performance in information extraction from scientific text **(Wang et al. 2024)**. However, no existing resource applies these tools to the scale of the full AMR literature while systematically linking the extracted evidence to the established reference databases **(Papp and Solymosi 2022)**.

Here, we introduce *ResLit*, an automated pipeline that (i) assembles a comprehensive corpus of AMR-relevant PubMed records through exhaustive controlled-vocabulary, substance-term, and free-text queries; (ii) screens abstracts for relevance using a BioMistral-7B few-shot classifier; (iii) retrieves full-text articles through a multi-tier downloading pipeline; (iv) extracts structured evidence on resistance genes and mutations using a Qwen3 model, followed by a second model pass that audits every extracted entity against the source text; and (v) organizes the results into a publicly accessible relational database in which each entry is cross-linked to CARD, ResFinder, and the NCBI Reference Gene Catalog and assigned an evidence tier reflecting its degree of corroboration. *ResLit* serves two complementary communities: i) microbiologists seeking to triage and summarize the literature for a gene, mutation, organism, or phenotype of interest, and ii) database curators requiring literature-backed evidence for validation workflows, including the review of existing entries whose support may warrant re-examination (**Figure 1**).

**Figure 1.**
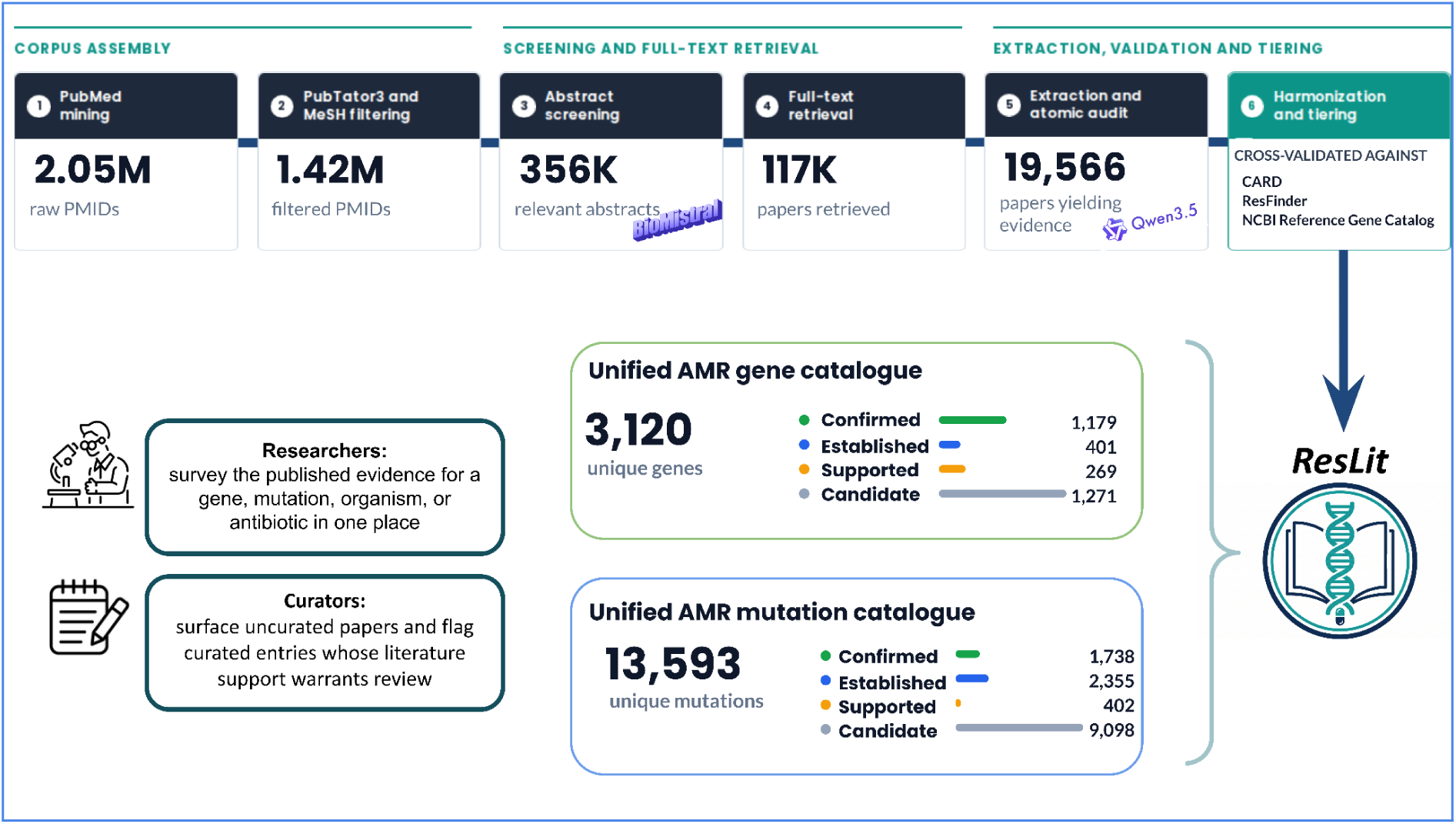
The ResLit pipeline. AMR-relevant PubMed records were assembled through 87 structured MeSH, substance-term, and free-text queries (2M unique PMIDs) and screened for relevance by BioMistral-7B. From the relevant papers, 117 K were retrieved in full text. The retrieved papers underwent a two-pass extraction with Qwen3-30B with deterministic post-processing. The extracted genes and mutations were matched to CARD, ResFinder, and the NCBI Reference Gene Catalog and assigned to one of four evidence tiers: confirmed (curated in one reference database and independently recovered by ResLit or curated in more than one reference database), established (curated in only one reference database without recovery by ResLit), supported (ResLit-extracted, ≥3 papers), and candidate (ResLit-extracted, <3 papers). The resulting catalog contained 3,120 genes and 13,593 mutations.

### Construction and content

#### Corpus assembly

##### Stage 1: eSearch retrieval

AMR-relevant PubMed records were retrieved via the NCBI Entrez eSearch API **(Sayers 2026)** using 87 structured queries (**Supplementary Section S1**) covering antibiotic classes, resistance mechanisms, resistance gene families, mobile genetic elements, and clinically relevant species. Queries combined MeSH descriptors and topical subheadings with pharmacological substance terms and free-text title/abstract searches for the named determinants and mechanisms. Free-text terms were added to reduce the risk of missing mechanism-focused reports that were incompletely captured by MeSH indexing. The search was restricted to publications dated between 1960 and 2025.

##### Stage 2: Metadata pre-filtering

The candidate corpus was reduced using a four-signal classifier with structured metadata. Entity annotations were drawn from PubTator3 **(Wei et al. 2024)**: the bulk gene, chemical, species, mutation, and relation files were indexed into a per-PMID table of Boolean flags recording whether each entity type was mentioned, together with a flag for high-value relation pairs (gene–chemical, variant–chemical, gene–disease, variant–disease). Bibliographic metadata (abstract, MeSH descriptors and subheadings, indexed substances, publication types, and language) were retrieved using the Entrez API and stored locally. A paper was considered to be relevant if any of the four signals were fired: a PubTator3 co-mention of a gene or mutation with a chemical; a curated AMR-mechanism MeSH descriptor co-occurring with an antibiotic or resistance-testing descriptor; a genetic-characterization MeSH subheading co-occurring with an AMR-associated substance or mechanism term; or a PubTator3 species mention together with both mechanism and drug MeSH descriptors (term lists in **Supplementary Section S2**). The signals are deliberately redundant so that incomplete annotation under any one indexing scheme does not exclude a genuinely relevant record. Passing records were then filtered to remove non-English papers and papers whose MeSH terms indicated a non-bacterial subject (fungal, viral, or parasitic) without any confirming bacterial terms. The classifier performance was evaluated against the ground-truth PMID set from the AmrProfiler database **(Skoulakis et al., 2025)**.

#### Screening and Full-text retrieval

##### Stage 3. Abstract Screening using Biomistral

Candidate records were screened for AMR relevance using BioMistral-7B. Screening was performed on the structured records already held in the local database: for each PMID, the classifier was presented with the article title, journal, abstract, MeSH descriptors, indexed substances, author keywords, and binary flags indicating whether the record mentioned a gene, chemical, or mutation. Where indexed substances matched a controlled list of AMR-associated terms, a single advisory line was appended, noting a possible resistance mechanism (**Supplementary Section S3**). Records lacking abstracts were excluded and not screened.

Classification was performed using a five-shot prompt. A system instruction defined AMR relevance broadly ( resistance genes and mutations, drug-target genes, gene cassettes, integrons, transposons, and other mobile genetic elements, efflux pumps, β-lactamases, and susceptibility studies) and stated explicitly that a paper qualifies even where the term “resistance” does not appear, so that cloning and characterization reports of resistance determinants are retained. All five examples were positive instances. This was a deliberate recall-oriented choice: the screening stage was tuned to admit false positives, which were subsequently eliminated during structured extraction, in preference to discarding relevant papers irrecoverably at this point. The classifier performance was evaluated against the ground-truth PMID set of the AmrProfiler database.

##### Stage 4. Full text retrieval

Full texts were retrieved through a multi-tier retrieval pipeline (**Supplementary Section S4**) that sequentially queried multiple open-access platforms and publisher TDM sources for each paper. First, article identifiers (PMID, PMCID, and DOI) were harmonized using the NCBI PMC ID Converter API to enable cross-platform retrieval (https://pmc.ncbi.nlm.nih.gov/tools/idconv/). Articles available in the PubMed Central (PMC) Open Access (OA) Subset were retrieved from PMC OA resources (https://europepmc.org/downloads), primarily using the OA bulk archive, PMC OA BioC **(Comeau et al. 2019)**, and Europe PMC **(Rosonovski et al. 2024)** full-text XML files. For papers with PMC identifiers but no corresponding OA Subset entry, the PMC eFetch XML API and the PMC web interface were used as alternative sources. For articles outside the PMC OA Subset, candidate open-access locations were identified through CrossRef, Unpaywall, which aggregate open-access availability across publishers and repositories (https://www.crossref.org/documentation/retrieve-metadata/rest-api/ , https://apis.io/providers/unpaywall/. Subscription-restricted papers were subsequently retrieved via the journal’s Text and Data Mining (TDM) service, accessed under the institution’s subscription agreement, including Elsevier, Wiley, Springer, subject to publisher-specific access policies and rate limits. Downloaded texts were stored as plain text, with HTML/XML tags removed and section structure preserved, where reliably parseable.

#### Extraction, Validation and Tiering

##### Stage 5: Structured information extraction

Structured AMR evidence was extracted from each retrieved full text using Qwen3-30B-A3B (https://huggingface.co/Qwen/Qwen3-30B-A3B) in two model passes, which were separated and followed by deterministic post-processing. Both passes applied unchanged to every document.

The first pass rendered each paper into a single JSON record of paper-level metadata and two entity collections: genes and mutations. The extraction prompt enforced three principles central to the pipeline’s precision-over-recall design: a results-only rule, admitting only entities arising from the paper’s experimental findings rather than its citations of other work; a validation gate, requiring a demonstrated resistance phenotype; and a separation rule directing genes that confer resistance only when mutated or inactivated (regulators, biosynthesis genes, and chromosomal core genes such as *gyrA*, *parC*, and *rpoB*) to the mutation collection. Papers exceeding the length threshold were chunked with overlap and merged.

A second pass audited every extracted entity individually against the source text, setting unsupported fields to null, and removing unsupported or misclassified entities. Deterministic post-processing was applied after the audit, handling of normalized gene names, filtered antibiotic assignments against a gene-family to-drug-class map, and discarded entries lacking literal support in the paper. Per-paper records were written individually, allowing the interrupted runs to resume. Full prompts, output schema, blocklists, drug-class map, and post-processing sequence are provided in **Supplementary Section S5**.

##### Stage 6: Harmonization and tiering

ResLit records were harmonized against curated reference data assembled in parallel from CARD, ResFinder, and the NCBI Reference Gene Catalog (NCBI RGC). These databases were downloaded on 01 May 2026. The three reference sources were reshaped into a common schema, their protein sequences pooled and clustered at 100% identity using CD-hit **(Li and Godzik 2006)**, and records that clustered together across sources were merged, so that a determinant reported by more than one database was represented once with its source memberships recorded. Gene and allele names in both the ResLit and reference catalogues were canonicalized against a shared gene-family reference, with β-lactamase alleles resolved to their bla-prefixed form. In the final alignment step, ResLit gene names were rewritten to the capitalization used in the reference catalogue, without adding, dropping, or otherwise altering records.

The Reslit catalogue was filtered to include entries carrying a recognized bacterial organism and a recognized antibiotic, with organism and drug names standardized against controlled vocabularies during the process. Mutation entries additionally underwent a nucleotide-versus-protein position consistency check, and promoter mutations identified by negative nucleotide coordinates were exempted from the requirement for a named antibiotic. Genes and mutations were then loaded into a relational database, together with a paper table derived by deduplicating PMIDs across both catalogues. A full description of the pipeline is provided in **Supplementary Section S6**.

At load time, the reference database source labels were normalized to a common set so that each ResLit determinant could be compared with its presence in CARD, ResFinder, and the NCBI RGC. Based on this comparison, every gene and mutation was assigned one of four evidence tiers defined by the intersection of two independent lines of evidence, presence in a curated reference database, and recovery by ResLit from the primary literature:

##### Confirmed

Corroborated by more than one independent line of evidence, either present in two or more reference databases or present in one reference database and independently recovered by ResLit.

##### Established

Curated in one reference database but not recovered by ResLit, either because the source publication lies outside the retrieved corpus or because extraction did not identify the association.

##### Supported

Absent from all three reference databases but reported in three or more independent studies in the ResLit corpus.

##### Candidate

Absent from all three reference databases and reported in fewer than three papers.

#### Database construction

ResLit is served by a web application built on Next.js 16, using server-rendered pages and Server Actions, and deployed as a single Docker container on Railway. Data are held in an embedded SQLite database; the database file resides on a persistent volume, with schema migrations applied automatically at deployment. Keeping the data local to the application container avoids the network round-trip and egress cost of a hosted service, which suits the read-heavy access pattern of public reference resources. Curator authentication is self-contained: passwords are hashed with scrypt, and sessions are opaque tokens stored as SHA-256 hashes. Interactive browsing over catalogues of this size is served through bounded, server-side queries, keeping the browse-page load times fast.

#### Database content

The Entrez eSearch strategy, comprising structured MeSH, substance-term, and free-text queries, retrieved 2,057,493 unique PMIDs spanning publications from 1960 to 2025. A four-signal metadata filter combining PubTator3 entity annotations with MeSH descriptors, subheadings, and indexed substances retained 1,543,369 candidate papers. Removal of non-English records (79,380) and records indexed to a non-bacterial subject (43,402) left 1,420,587 papers.

Pipeline recall was benchmarked at each stage against the AmrProfiler ground-truth set (2,628 gene-associated and 365 mutation-associated reference PMIDs). The metadata pre-filter retained 91.1% of gene-associated (2,394/2,628) and 97.8% of mutation-associated (357/365) reference PMIDs, confirming that the search preserved the majority of known AMR literature.

Next, BioMistral-7B few-shot screening classified 356,262 of the retained records as AMR-relevant, recovering 86.9% of gene-associated (2,285/2,628) and 97.3% of mutation-associated (355/365) reference PMIDs against the same set. Full texts were successfully retrieved for 117,112 of the screened papers and were carried forward for structured extraction. Because extraction operates on full text only, ground-truth recall at this point fell to 61.5% for gene-associated (1,617/2,628) and 65.5% for mutation-associated (239/365) reference PMIDs, reflecting papers behind paywalls or otherwise unavailable for download. Determinants described exclusively in such papers were not lost from the database. If they are curated in CARD, ResFinder, or the NCBI RGC, they enter ResLit through the reference pipeline and are retained in the established tier. Thus, the loss of full-text access affects ResLit’s independent literature recovery of a determinant but does not eliminate the determinant from the database.

Structured extraction using QWEN-3 yielded verified AMR evidence from 19,566 papers. The released database contains 3,120 unique resistance genes across 54,175 gene-paper records and 13,593 unique mutations across 26,094 mutation-paper records, each of which is cross-referenced against CARD, ResFinder, and the NCBI Reference Gene Catalog.

##### Gene coverage across the databases

Pooling the four resources yielded 3,120 unique resistance genes, of which only 480 (15%) were present in all four and 1,941 (62%) in just one, reflecting a substantial disagreement among the curated references. Against this fragmented landscape, ResLit was the largest single catalogue, with 2,546 genes, and recovered a substantial share of each reference database: 885 of 1,272 CARD genes (70%), 840 of 1,138 NCBI RGC genes (74%), and 537 of 638 ResFinder genes (84%), showing that its coverage is distributed across all three resources rather than concentrated in any one (**Figure 2**). A further 1,540 genes were reported in the primary literature but were absent from all three reference databases, which was the single largest contribution to the pool, and the entries were the most directly useful as candidates for curation.

**Figure 2.**
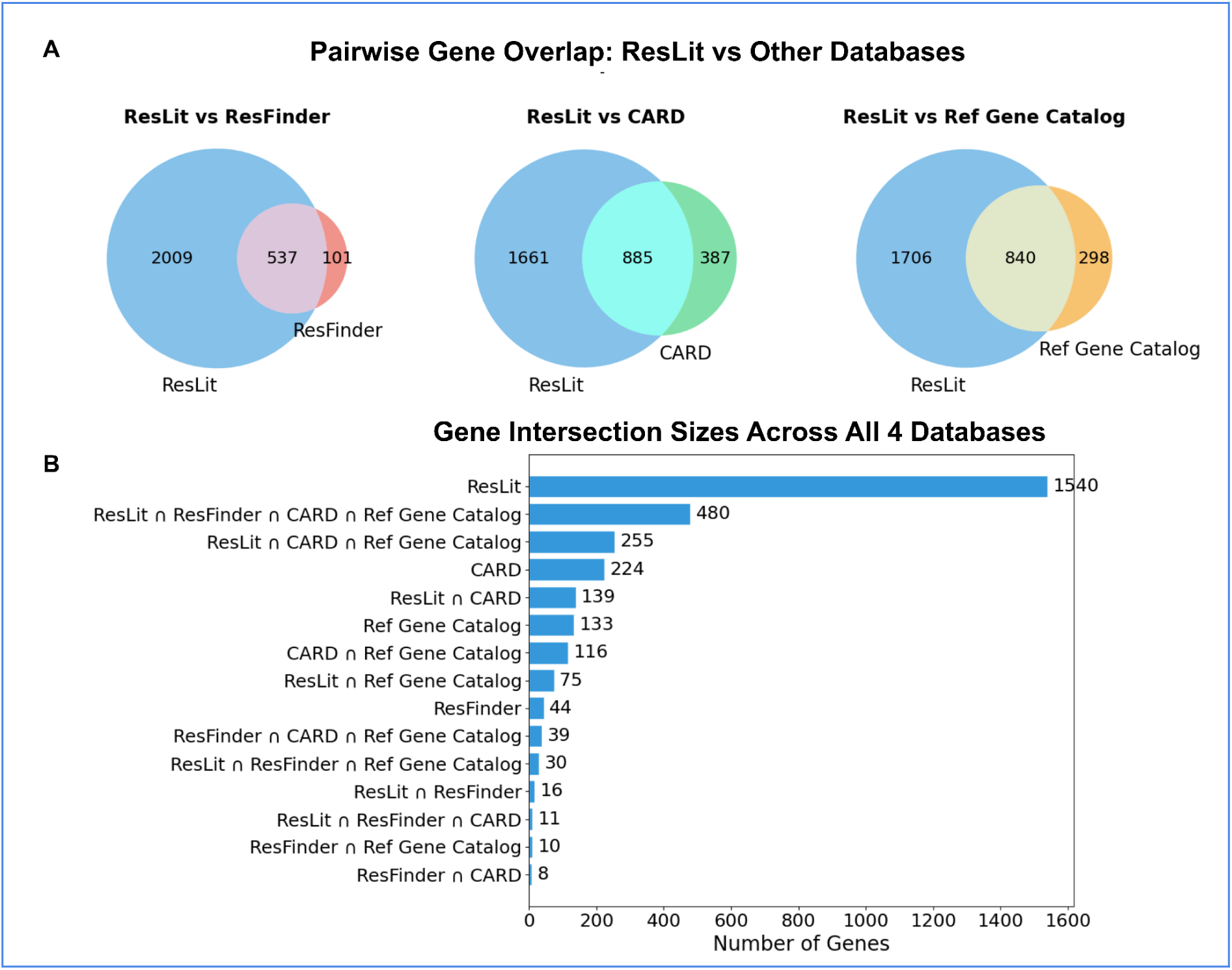
Overlap between the ResLit gene catalogue and CARD, ResFinder, and NCBI Reference Gene Catalog reference databases. **(A)** Pairwise overlap between ResLit and each reference database, showing genes unique to each catalogue and those that are shared. ResLit recovered 537 of the 638 ResFinder genes, 885 of the 1,272 CARD genes, and 840 of the 1,138 Reference Gene Catalog genes, while contributing a large set of genes (2,009, 1,661, and 1,706 in the three comparisons) that were not present in each. **(B)** Sizes of all 15 intersections across the four catalogues, ordered by size. ResLit contributed 1,540 genes absent from all three curated references, the single largest category, whereas 480 genes were shared by all four. Gene names were normalized to a common form before comparison.

##### Mutation coverage across the databases

The mutation catalogues showed the same fragmentation as the genes; a lower overlap is expected, given the greater difficulty in matching mutation notation across resources. ResLit was by far the largest catalogue, with 10,939 unique mutations, compared to 2,283 in CARD, 1,424 in NCBI RGC, and 1,236 in ResFinder. ResLit recovered 838 of 2,283 CARD mutations (37%), 651 of 1,424 NCBI RGC mutations (46%), and 490 of 1,236 ResFinder mutations (40%) (**Figure 3**). Only 69 mutations were shared among the four catalogues. The 9,500 mutations discovered by ResLit were reported in the primary literature but were absent from all three reference databases, underscoring the extent of uncurated mutation-level resistance literature.

**Figure 3.**
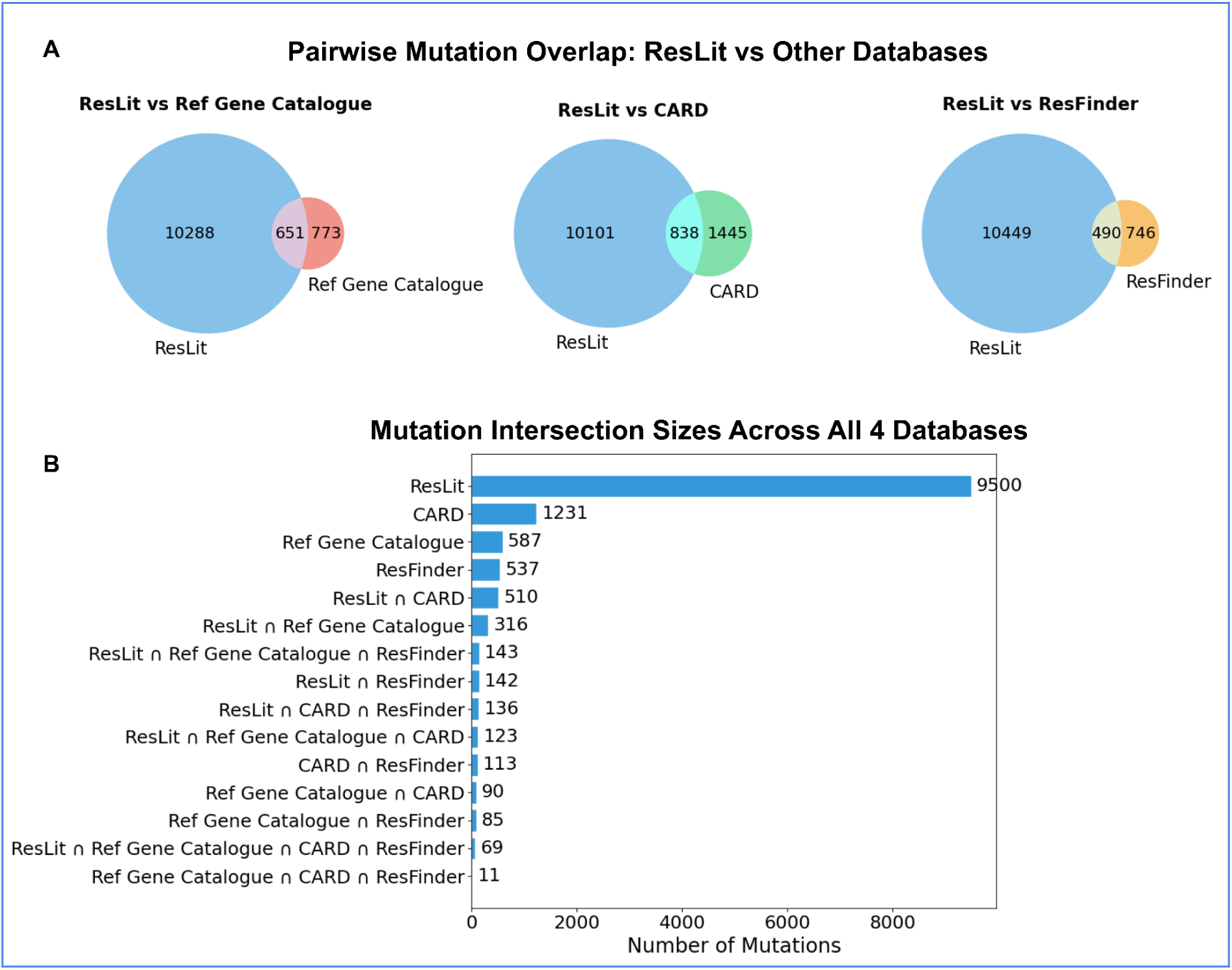
Overlap between the ResLit mutation catalogue and the CARD, NCBI Reference Gene Catalog, and ResFinder reference databases. **(A)** Pairwise overlap between ResLit and each reference database, showing unique mutations for each catalogue and those that are shared. ResLit recovered 651 of 1,424 NCBI RGC mutations, 838 of 2,283 CARD mutations, and 490 of 1,236 ResFinder mutations, while contributing a large set of mutations (10,288, 10,101, and 10,449 in the three comparisons) not present in each. **(B)** Sizes of all mutation intersections across the four catalogues, ordered by size. ResLit contributes 9,500 mutations absent from all three curated references, the single largest category, whereas 69 mutations are shared by all four. Mutations were matched to gene identity together with the reported change.

##### Evidence tiers

The cross-database comparisons above distinguish the determinants by which resources contain them; ResLit combines this with literature evidence to assign each gene and mutation to one of four evidence tiers **(Table 1**). The four evidence tiers are as follows: confirmed (curated in one reference database and independently recovered by ResLit or curated in more than one reference database), established (curated in only one reference database without recovery by ResLit), supported (ResLit-extracted, ≥3 papers), and candidate (ResLit-extracted, <3 papers). Of the 1,580 genes and 4,093 mutations curated in at least one reference database, ResLit independently recovered 1,179 genes (74.6%) and 1,738 mutations (42.5%) from the literature, with the lower mutation recall reflecting the greater difficulty of extracting mutation notation from free text. The remaining 1,540 genes and 9,500 mutations were reported in the literature but were absent from all three reference databases and constitute ResLit’s principal contribution to curation: determinants with published support that no curated resource records.

**Table 1.** Evidence-tier distribution of ResLit genes and mutations.

| **Tier** | **Genes** | **Mutations** |
| --- | --- | --- |
| Confirmed | 1,179 | 1,738 |
| Established | 401 | 2,355 |
| Supported | 269 | 402 |
| Candidate | 1,271 | 9,098 |
| **Total** | **3,120** | **13,593** |

## Utility and discussion

### Database interface and accessibility

The ResLit web portal provides search and browse access to all database entries through two parallel catalogues, one for resistance genes (**Figure 4**) and one for mutations (**Figure 5**), along with a dedicated curation interface (**Figure 6**).

**Figure 4.**
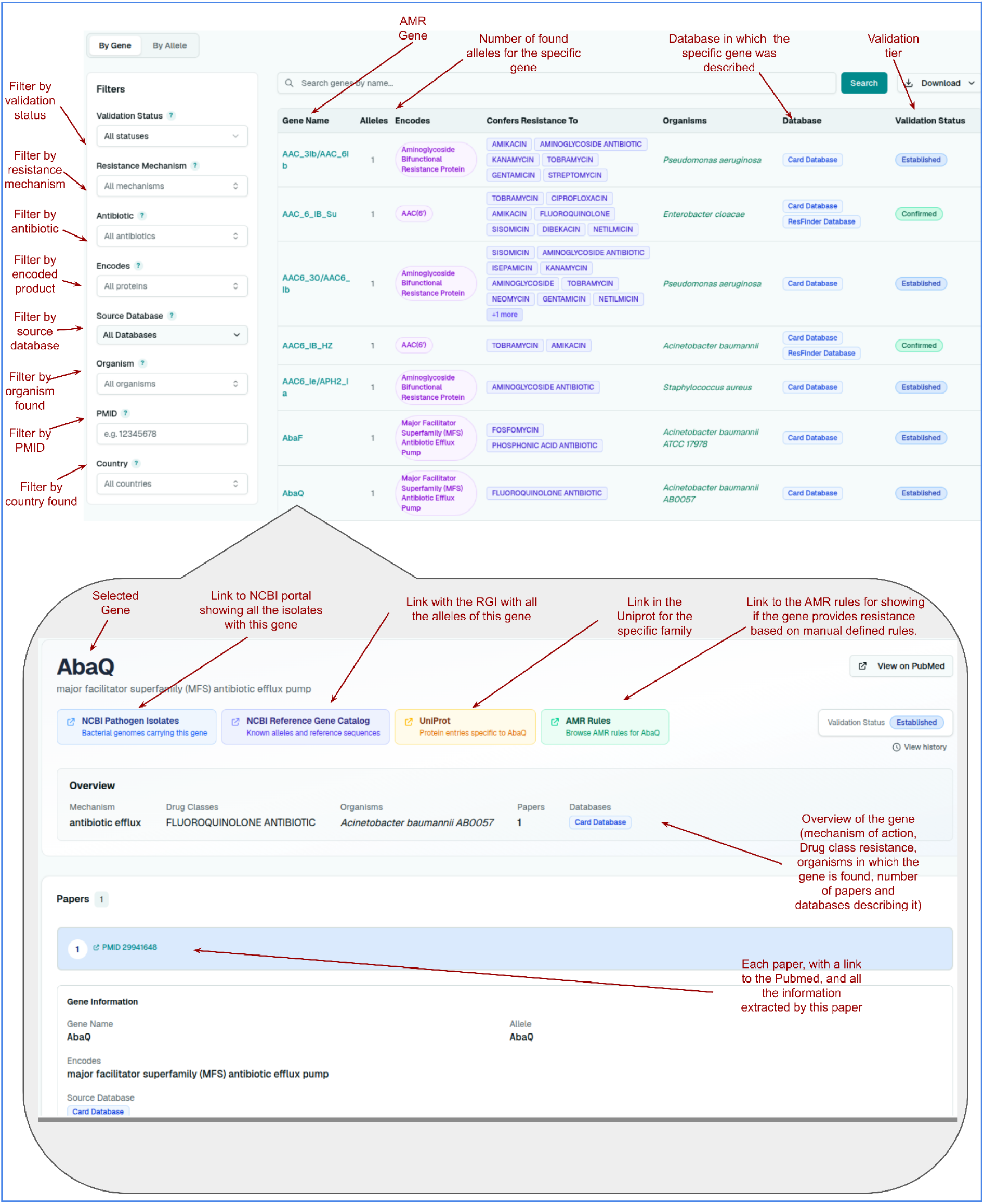
The ResLit Gene Catalogue. Browse interface for resistance genes. The faceted sidebar (left) filters by validation status, resistance mechanism, antibiotic, encoded product, source database, organism, PMID, and country; the central table lists each gene with its alleles, encoded product, antibiotics to which it confers resistance, host organisms, source database, and evidence tier, and supports free-text search and tabular download. Selecting a gene opens a detailed panel (bottom) providing its mechanism, drug classes, organisms, allele designation, sequence and protein accessions, and contributing publications, with links to NCBI Pathogen Isolates, the NCBI Reference Gene Catalog, UniProt, and AMR Rules, and a curation-history view.

**Figure 5.**
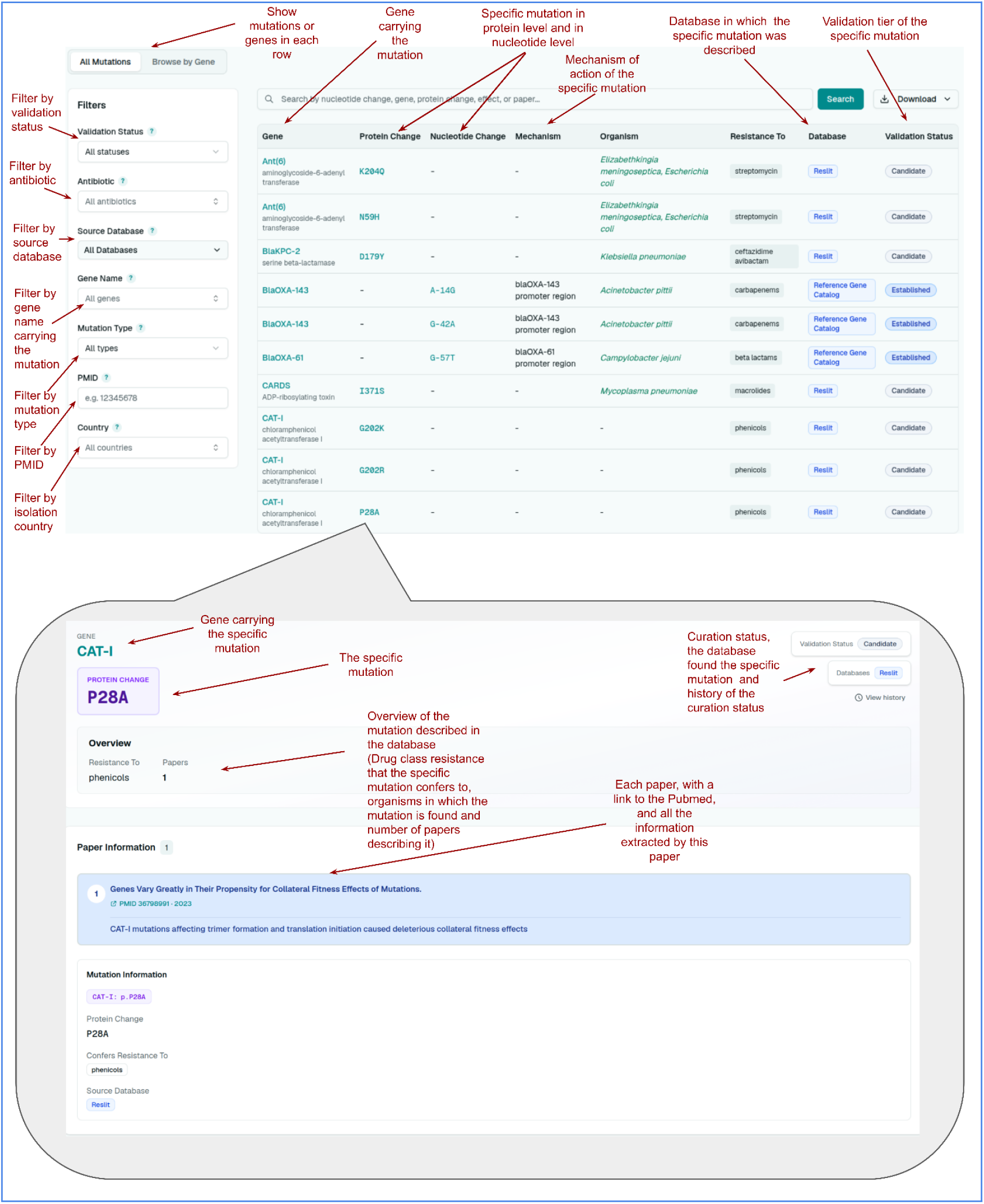
The ResLit mutation catalogue. Browse interface for resistance mutations, filterable by validation status, antibiotic, source database, gene name, mutation type, PMID, and country, and browsable as a flat list or grouped by target gene. The table lists each mutation with its gene, protein- and nucleotide-level changes, mechanism, host organism, antibiotics affected, source database, and evidence tier. The detailed panel (bottom) presents the changes, organisms, antibiotics, validation methods, and source databases, together with each supporting publication and the evidence statement extracted from it.

**Figure 6.**
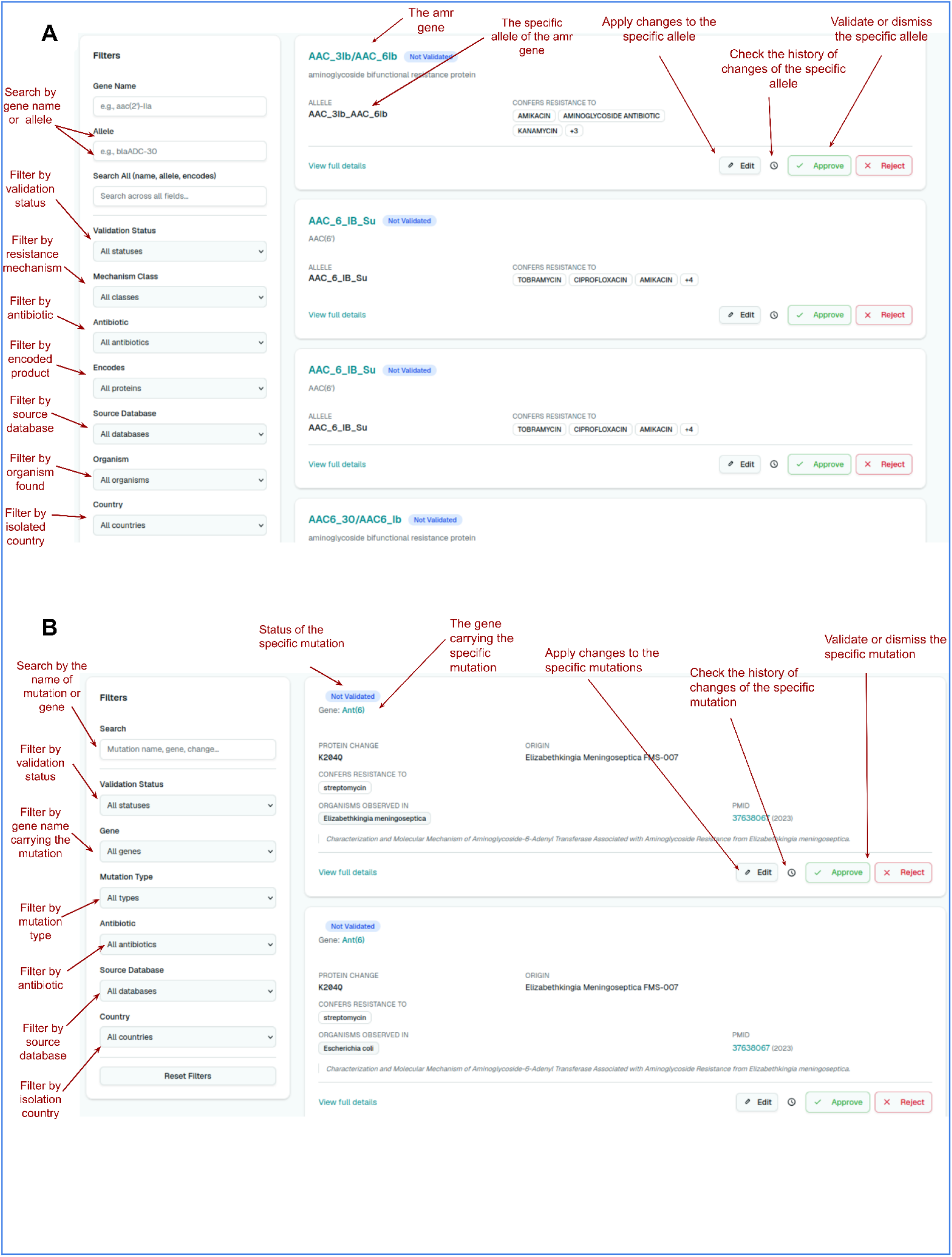
The curator interface. Authenticated curators reviewed genes and mutations using parallel curation views. Each entry is shown with its extracted fields and validation status and carries per-entry controls to edit the record, view its revision history, and approve or reject it, along with a link to the corresponding browse record. The same filters as the public catalogues allow curators to target entries based on validation status, source database, organism, and other criteria. (A) Gene curation view, showing each resistance gene with its allele, encoded product, and antibiotics to which it confers resistance. (B) Mutation curation view showing each mutation with its target gene, protein change, origin, and supporting publication.

Both catalogues present a filterable table alongside faceted sidebars. Genes may be filtered by validation status, resistance mechanism, antibiotic, encoded product, source database, organism, PubMed identifier, and country of origin of the isolate; mutations by validation status, antibiotic, source database, gene name, mutation type, PMID, and country of origin of the isolate. Free-text search operates over gene names in the gene catalogue and gene name, protein change, and nucleotide change in the mutation catalogue. Mutations can also be browsed by their target genes. Every row displays its evidence tier and source databases, so that the provenance of an entry is visible without opening it.

Selecting an entry opens a detailed panel. For genes, the panel presents the encoded product and mechanism, drug classes, organisms tested, allele designation, sequence and protein accessions, contributing publications with PMID and year, and outbound links to the corresponding NCBI Reference Gene Catalog, NCBI Pathogen Isolates, UniProt, and AMR Rules records (**Figure 4**). For mutations, it presents the protein or nucleotide change, position within the molecule, organisms observed, antibiotics affected, validation method, and supporting publications with the extracted evidence statement for each (**Figure 5**). Both panels expose a revision history, allowing any entry to be traced through subsequent curatorial amendments to the database.

Signed-in curators access a dedicated review interface (**Figure 6**), in which each gene or mutation is presented with its extracted fields and per-entry controls to edit, approve, or reject it. Curators can correct a field, flag an entry as unsupported by its cited paper, or reassign it between the gene and mutation catalogues, and can move directly from a curation entry to its corresponding browse record. Amendments are recorded as revisions against the original machine-extracted value rather than replacing it, so that the extraction pipeline remains auditable against expert judgement and every version of an entry remains citable. Candidate-tier entries, which are absent from all reference databases, can be isolated by filtering and are the natural starting point for curation.

The full contents of either catalogue are available for download in tabular format, with the active filter set applied, so that a query (for instance, all candidate-tier mutations in a given organism) can be exported directly as a working list rather than being reconstructed offline.

### Surfacing candidate discrepancies in reference databases

To further illustrate the use of ResLit, we examined entries in CARD **(McArthur et al., 2013)**, ResFinder **(Bortolaia et al., 2020)**, and the NCBI RGC **(Goldfarb et al., 2025),** for which ResLit’s search of the primary literature returned no supporting evidence (Established tier). A missing recovery need not indicate an error: the cited publication may be paywalled and therefore outside the ResLit corpus or may not state the association in a form the extraction model could identify. In a subset of cases, however, inspection of the cited source reveals a genuine discrepancy warranting curator review.

Table 2. Representative candidate discrepancies were identified by literature cross-referencing. The source publications examined included **(Zimenkov et al. 2013; Zhang et al. 2020; Rice et al. 2000; Springer et al. 2001; Nessar et al. 2011; Suzuki et al. 1998; Zhang et al. 2013)** the corresponding PMIDs and the basis for review are provided for each entry.

**Table 2.** presents representative examples, all of which surfaced during routine cross-referencing rather than targeted searches. In each case, an entry curated with a specific resistance mutation traces to a paper that does not support that mutation as stated.

| Source | Entry | Organism | Drug | PMID | Basis for review |
| --- | --- | --- | --- | --- | --- |
| ResFinder | *eis* promoter G-10A | *M. tuberculosis* | kanamycin | 23995137 | Cited paper does not discuss this variant |
| ResFinder | *eis* A-13G | *M. tuberculosis* | kanamycin | 23705640 | Cited paper does not discuss this variant |
| NCBI RGC | blaOXA-143 A-14G | *A. pittii* | carbapenem | 32582088 | Cited paper reports variants of blaOXA-499, not blaOXA-143 |
| NCBI<br>RGC | blaSHV<br>C-112A | *K.*<br>*pneumoniae* | cefiderocol /<br>cephalosporin | 10639363 | Cited paper<br>does not<br>discuss this<br>variant |
| CARD | *rrs* a503c | *M.*<br>*smegmatis* | streptomycin | 11557484 | Cited paper<br>does not<br>discuss this<br>variant |
| CARD | *rrs* c1376t | *M.*<br>*abscessus* | amikacin,<br>gentamicin,<br>kanamycin,<br>tobramycin | 21652621 | Cited paper<br>does not<br>discuss this<br>variant |
| CARD | *rrs* c1402a | *M.*<br>*tuberculosis* | kanamycin | 9574680 | Cited paper<br>does not<br>appear to<br>establish that<br>this variant<br>confers<br>resistance |

Three patterns accounted for most cases. The first is allele misattribution: for blaOXA-143, the cited publication characterized variants of blaOXA-499, and the substitution was recorded against a different allele of the same family. The second is coordinate-system ambiguity: without a stated reference sequence, one reported mutation can be numbered under several schemes, generating apparently distinct entries for a single observation. The third is over-attribution: a paper reports observing a variant but does not present it as resistance-conferring, and the entry nonetheless records it as such.

These are candidate discrepancies raised for expert review, not confirmed errors; several turn on interpreting whether a paper’s evidence establishes causation rather than a demonstrable mismatch. They also point to a broader curation need: for mutation entries, in particular, recording the reference genome or sequence against which the coordinates are defined would remove much of the ambiguity that produces these cases. Their value here is in showing that ResLit can direct curator attention to specific entries and papers at a scale that manual review cannot reach.

## Discussion

*ResLit* addresses a gap in the AMR informatics landscape: the absence of a large-scale, automated, literature-linked resource that extracts and organizes AMR evidence at a resolution required to support both hypothesis-driven research and database curation. Existing AMR databases are invaluable but rely on expert curation that cannot keep pace with the growth of primary literature. *ResLit* complements these tools by operating at scale, providing an updatable evidence layer indexed against CARD **(McArthur et al., 2013)**, ResFinder **(Bortolaia et al., 2020)**, and the NCBI RGC **(Goldfarb et al., 2025)**.

The staged architecture, BioMistral-7B for relevance screening **(Labrak et al. 2024)**, Qwen3-30B for structured extraction, and a second Qwen3 pass auditing every extracted entity against its source text **(Yang et al. 2025)**, reflects a deliberate allocation of recall and precision to different points in the pipeline. Screening is tuned for recall, admitting false positives that structured extraction subsequently discards by failing to produce valid outputs for out-of-topic papers. Extraction and auditing are tuned for precision, on the reasoning that at this scale, a small false-positive rate produces thousands of incorrect entries and decreases the curator’s trust inthe resource on which they depend, whereas a missed determinant is usually recoverable from another paper reporting it. Running the smaller screening model over the full corpus and reserving the larger extraction model for the screened subset was also substantially cheaper than applying the extraction directly to all candidates.

The four-tier evidence scheme makes the limitations of the pipeline legible rather than concealing them. The Established tier (determinants curated in a reference database that ResLit did not recover) is a direct measure of the coverage gap, and the disparity between genes (75% recovery) and mutations (43%) locates that gap precisely. Crucially, this gap reflects incomplete *recovery*, not lost data: curated mutations ResLit did not independently recover are still present in ResLit and retained in the Established tier through the reference pipeline.

Several limitations warrant acknowledgement. Extraction accuracy depends on the model’s fidelity to scientific full texts, which varies with writing style and the specificity of resistance reporting. This was most pronounced for mutations, where recovery reached only 43%. Mutations are intrinsically harder to extract than genes: notation varies between authors, and a position number is often ambiguous without a stated reference sequence; therefore, the same change can be written in several ways depending on the genome or protein it is numbered against. This is most acute for rRNA genes, such as *rrs* and *rrl*, numbered under competing organism-native and *E. coli* conventions. The same ambiguity that lowers ResLit’s recovery also leads curated databases to record one real mutation as several apparently distinct entries.

Insertions and deletions are under-represented relative to substitutions: substitutions carry a compact, standardized notation that the model can transcribe reliably, whereas indels are typically described in prose, as a frameshift, truncation, or named deletion without stated coordinates, which the model annotates less reliably. Because extraction operates on full text, coverage is also bounded by access: many relevant papers are behind paywalls and could not be retrieved, and because paywalled titles cluster by publisher, the resulting gap is systematic rather than random. Given the public health stakes of antimicrobial resistance, broader open access to this literature would materially improve the completeness of resources such as ResLit. Coverage was further limited to English-language, PubMed-indexed literature, excluding regional and non-indexed journals. Finally, cross-referencing to reference databases relies on name matching; therefore, a proportion of candidate-tier entries may represent variant names for previously characterized determinants rather than genuinely novel ones, a possibility that the tier label is intended to signal rather than obscure.

The discrepancies in Table 2 point to a use for *ResLit* beyond gene and mutation search or discovery of related AMR publications. Because every entry retains a link to its source publication, the resource can direct curator attention not only to uncurated determinants but to curated entries whose supporting literature may warrant re-review. Both the allele misattributions and the coordinate-system ambiguities we observed are the kind of error that propagates once an accession enters downstream tooling, and both are difficult to detect by manual review at scale. We raise these cases as candidates for expert adjudication, not as confirmed errors, and we regard the ability to generate such candidates systematically as one of the resource’s more useful contributions.

*ResLit* is intended to function as a continuously curated resource, rather than a static release. Authenticated curators can amend any entry directly, with corrections recorded as revisions against the original machine-extracted value, so that expert judgement accumulates in the database while the pipeline output remains auditable against it. The database will be re-run against new literature and re-indexed against each major release of the reference databases on an annual basis. By providing open, bulk-downloadable, machine-readable AMR evidence at scale, together with a mechanism for expert correction, *ResLit* aims to accelerate the translation of primary research into updated surveillance databases, improved predictive models and evidence-based clinical guidelines.

## Data Availability Statement

The ResLit database is freely available at https://www.reslit.info without registration. Only curators need to register in order to be able to edit, validate or reject entries. The complete gene and mutation catalogues can be downloaded in tabular (TSV) format directly from the web interface, with or without filters. The pipeline source code, corpus assembly, metadata pre-filtering, relevance screening, structured extraction, and harmonisation, is available at https://github.com/Georgakopoulos-Soares-lab/ResLit. The ground-truth PMID sets and reference database snapshots used to benchmark recall originated from the AmrProfiler database.

## Acknowledgements

This work was supported by startup funding provided to Ilias Georgakopoulos-Soares by the University of Texas, Austin. G.A.P. and A.G. were supported by the Hellenic Foundation for Research and Innovation (H.F.R.I.) under the “Third Call for H.F.R.I. Research Projects to support faculty members and researchers” [23592-EMISSION].

## Author Contributions

A.S., H.X., and I.G.-S. wrote the manuscript. A.S. and H.X. created the pipeline for finding and downloading papers. A.S. created the LLM extraction pipeline. A.S., A.G., and G.P. harmonized the genes and mutations. A.S. and K.A.P. developed the online tools. I.G.-S. supervised the work. All the authors have reviewed the manuscript.

