## Supplementary material for "ResLit: A Large-Scale Automated Literature Mining Database for Antimicrobial Resistance"

**ResLit: Supplementary Information**

This document provides the supplementary methods underlying the ResLit pipeline. It is organised to follow the pipeline in order, one section per stage; section numbers match the pipeline stage numbers used in the main text and in Figure 1.

**Contents**

**Section S1**	Stage 1 : corpus assembly, Entrez search strategy and query panel

**Section S2**	Stage 2 : metadata pre-filter: signals and term lists

**Section S3** Stage 3 : relevance screening: classifier prompt, examples, and substance terms

**Section S4**	Stage 4 : full-text retrieval

**Section S5** Stage 5 : structured extraction, model parameters, schema, filters, and prompts

**Section S6** Stage 6 : harmonization and tiering, reference assembly, post-processing chains, and tier logic

**Section S1. Corpus assembly** - **Stage 1: Entrez retrieval**

This section documents how the ResLit candidate corpus was assembled: the search strategy issued against PubMed, the complete panel of queries with the number of records each returned, and the deduplication that produced the final candidate set.

**S1.1 Search strategy**

AMR-relevant PubMed records were retrieved through the NCBI Entrez eSearch API using a panel of 87 structured queries covering antibiotic classes, resistance mechanisms, resistance gene families, mobile genetic elements, and clinically relevant organisms. The panel combined three complementary query types to maximise coverage of both well-indexed and sparsely indexed records: Medical Subject Heading (MeSH) descriptors paired with topical subheadings (principally genetics, drug effects, pharmacology, and biosynthesis); pharmacological substance terms; and free-text title/abstract searches targeting named determinants and mechanisms. The free-text component was included specifically to capture mechanism-focused reports whose MeSH indexing is incomplete or absent, which controlled-vocabulary queries alone systematically miss.

Each query was restricted to publications dated 1960-2025. Because the Entrez API caps any single query at 9,999 returned records, queries were executed under a recursive date-windowing strategy: a query was first issued for each calendar year, and any year-level query exceeding the cap was automatically subdivided by month, then by day, until every sub-query returned a tractable number of records. This guaranteed complete retrieval of high-yield queries, the broadest returned in excess of 500,000 records each. Requests were rate-limited to a minimum inter-request interval of 0.35 s in line with NCBI usage policy, and wrapped in an exponential-backoff retry handler (up to seven attempts) covering HTTP 429 rate-limit responses, HTTP 500 server errors, and transient connection failures, so that no query was dropped as a result of intermittent service interruption.

PubMed identifiers (PMIDs) returned by all queries were pooled and deduplicated by set union, yielding 2,057,493 unique candidate PMIDs. Total retrieval time was approximately 87 minutes.

*Zero-yield queries.* Queries 6 and 15 returned no records because bacterial and resistance are not valid MeSH descriptors. Both are retained in the panel below for completeness of the executed strategy. Neither affects corpus coverage: each is a superset of another query in the panel that did return records (queries 7 and 16 respectively), so no records were lost.

**S1.2 Query panel**

The 87 queries are listed below in execution order, grouped by target category. Each is annotated with the number of unique PMIDs it returned. Query syntax is reproduced exactly as issued to the Entrez API.

**1. General antimicrobial resistance**	(Q1-Q5)

*Broad entry-point queries combining resistance concepts with gene or mutation terms.*

**Q1**	-	84,927 PMIDs

("drug resistance, microbial"[MeSH Terms] OR "antimicrobial resistance"[All Fields]) AND (gene[All Fields] OR mutation[All Fields])

**Q2**	-	47,022 PMIDs

("antibiotic resistance"[Title/Abstract] OR "multidrug resistance"[Title/Abstract]) AND (gene OR mutation)

**Q3**	-	36,883 PMIDs

"drug resistance, microbial"[MeSH Terms] AND ("genes"[MeSH Terms] OR "mutation"[MeSH Terms])

**Q4**	-	55,587 PMIDs

"antimicrobial resistance"[Title/Abstract]

**Q5**	-	310,982 PMIDs

(("resistance"[MeSH Terms] OR "resistance"[All Fields]) AND ("bacteria"[MeSH Terms] OR "bacteria"[All Fields] OR "bacterial"[All Fields]))

**2. MeSH descriptors with topical subheadings**	(Q6-Q17)

*Descriptor-subheading pairs targeting the genetics of resistance determinants, their carriers, and their bacterial hosts.*

**Q6**	-	0 PMIDs

(bacterial[mh] AND genetics[sh])

**Q7**	-	531,169 PMIDs

(bacteria[mh] AND genetics[sh])

**Q8**	-	104,222 PMIDs

(Plasmids[mh] AND genetics[sh])

**Q9**	-	18,911 PMIDs

(beta-Lactamases[mh] AND genetics[sh])

**Q10**	-	216 PMIDs

(Cephalosporinase[mh] AND genetics[sh])

**Q11**	-	201,834 PMIDs

("Bacterial Proteins"[mh] AND genetics[sh])

**Q12**	-	207,240 PMIDs

(Enterobacteriaceae[mh] AND genetics[sh])

**Q13**	-	24,222 PMIDs

(Salmonella[mh] AND genetics[sh])

**Q14**	-	54,031 PMIDs

"Gene Expression Regulation, Bacterial"[mh]

**Q15**	-	0 PMIDs

(resistance[mh] AND genetics[mh])

**Q16**	-	163,719 PMIDs

("Drug Resistance"[mh] AND genetics[sh])

**Q17**	-	75,839 PMIDs

("Anti-Bacterial Agents"[mh] AND genetics[sh])

**3. Broad MeSH combinations**	(Q18-Q24)

*High-yield descriptor combinations providing baseline coverage of the mutation and resistance literature.*

**Q18**	-	775,872 PMIDs

Mutation[mh]

**Q19**	-	140,053 PMIDs

("Drug Resistance, Microbial"[mh] AND (bacterial[mh] OR bacteria[mh] OR genetics[mh]))

**Q20**	-	48,004 PMIDs

("Drug Resistance, Microbial"[mh] AND (Bacillus[mh] OR Staphylococcus[mh] OR Escherichia[mh]))

**Q21**	-	39,474 PMIDs

((bacterial[mh] OR bacteria[mh]) AND genetics[mh])

**Q22**	-	23,442 PMIDs

(Mutation[mh] AND Plasmids[mh])

**Q23**	-	11,843 PMIDs

("Anti-Bacterial Agents"[mh] AND Mutation[mh])

**Q24**	-	24,636 PMIDs

("Drug Resistance, Microbial"[mh] AND Mutation[mh])

**4. β-lactams and β-lactamases**	(Q25-Q32)

*Cephalosporins, carbapenems, β-lactamase inhibitors, and β-lactamase biosynthesis and physiology.*

**Q25**	-	13,101 PMIDs

("Cephalosporins"[mh] OR "Cephalosporin Resistance"[mh]) AND ("Drug Resistance, Microbial"[mh] OR "beta-Lactamases"[mh])

**Q26**	-	302 PMIDs

(Cefpodoxime[mh] OR Ceftizoxime[mh]) AND ("beta-Lactamases"[mh] OR "Drug Resistance, Microbial"[mh])

**Q27**	-	2,637 PMIDs

("beta-Lactamase Inhibitors"[mh]) AND ("beta-Lactamases"[mh] OR "Drug Resistance, Microbial"[mh])

**Q28**	-	5,256 PMIDs

("beta-Lactamases"[mh] AND biosynthesis[sh]) AND ("bacteria"[mh] OR "Enterobacteriaceae"[mh])

**Q29**	-	9,907 PMIDs

("Klebsiella pneumoniae"[mh] OR "Klebsiella Infections"[mh]) AND ("beta-Lactamases"[mh] OR "Drug Resistance, Microbial"[mh])

**Q30**	-	108,178 PMIDs

("Anti-Bacterial Agents"[mh] AND ("Drug Resistance, Microbial"[mh] OR "beta-Lactamases"[mh]))

**Q31**	-	10,008 PMIDs

("Carbapenems"[mh] OR "Carbapenem Resistance"[mh]) AND ("beta-Lactamases"[mh] OR "Drug Resistance, Microbial"[mh])

**Q32**	-	29,595 PMIDs

("beta-Lactamases"[mh] AND (biosynthesis[sh] OR physiology[sh] OR isolation[sh]))

**5. Aminoglycosides and fluoroquinolones**	(Q33-Q34)

**Q33**	-	13,107 PMIDs

("Aminoglycosides"[mh] OR "Aminoglycoside Resistance"[mh]) AND ("Drug Resistance, Microbial"[mh])

**Q34**	-	10,008 PMIDs

("Fluoroquinolones"[mh] OR "Quinolone Resistance"[mh]) AND ("Drug Resistance, Microbial"[mh])

**6. Free-text searches for resistance determinants and mechanisms**	(Q35-Q42)

*Title/abstract queries targeting named determinants and mechanism vocabulary, included to capture mechanism-focused reports whose MeSH indexing is incomplete or absent.*

**Q35**	-	12,085 PMIDs

(("target site mutation" OR "target mutation" OR "point mutation")[Title/Abstract] AND ("antibiotic" OR "drug" OR "resistance")[Title/Abstract])

**Q36**	-	11,839 PMIDs

(("gyrA" OR "parC" OR "gyrB" OR "parE" OR "23S" OR "rpoB" OR "rpoC" OR "fusA" OR "ftsZ" OR "murA" OR "murB" OR "walK" OR "walR" OR "pbp")[Title/Abstract] AND ("mutation" OR "polymorphism" OR "variant")[Title/Abstract])

**Q37**	-	24,221 PMIDs

(("DNA gyrase" OR "topoisomerase" OR "RNA polymerase" OR "ribosomal protein" OR "penicillin binding protein" OR "murein biosynthesis")[Title/Abstract] AND ("mutation" OR "resistance")[Title/Abstract])

**Q38**	-	6,600 PMIDs

(("resistance mechanism" OR "mechanism of resistance" OR "resistance determinant")[Title/Abstract] AND (bacteria[All Fields] OR organism[All Fields] OR "species"[All Fields]))

**Q39**	-	6,328 PMIDs

((mutation[Title/Abstract] OR "amino acid substitution"[Title/Abstract]) AND (Salmonella[Title/Abstract] OR "Escherichia coli"[Title/Abstract] OR Staphylococcus[Title/Abstract] OR Pseudomonas[Title/Abstract] OR Acinetobacter[Title/Abstract] OR Klebsiella[Title/Abstract] OR Streptococcus[Title/Abstract]) AND (resistance[Title/Abstract] OR susceptibility[Title/Abstract]))

**Q40**	-	9,862 PMIDs

(("minimum inhibitory concentration" OR "MIC" OR "antimicrobial susceptibility")[Title/Abstract] AND (mutation[Title/Abstract] OR gene[Title/Abstract]) AND (resistance[Title/Abstract] OR "determination region"[Title/Abstract]))

**Q41**	-	49,615 PMIDs

(("beta-lactamase" OR "aminoglycoside modifying" OR "efflux" OR "acetyltransferase" OR "phosphotransferase")[Title/Abstract] AND (mutation[Title/Abstract] OR variant[Title/Abstract] OR resistance[Title/Abstract]))

**Q42**	-	49,543 PMIDs

(("gene expression" OR "upregulation" OR "downregulation" OR "overexpression" OR "regulatory")[Title/Abstract] AND ("resistance" OR "antibiotic")[Title/Abstract] AND (bacteria[All Fields] OR organism[All Fields]))

**7. Priority pathogens**	(Q43-Q44)

**Q43**	-	10,268 PMIDs

(Pseudomonas aeruginosa[mh] AND ("Drug Resistance, Microbial"[mh] OR "beta-Lactamases"[mh]))

**Q44**	-	5,331 PMIDs

(Acinetobacter baumannii[mh] AND ("Drug Resistance, Microbial"[mh] OR "beta-Lactamases"[mh]))

**8. Additional classes, mobile genetic elements, and ESBL/AmpC**	(Q45-Q49)

**Q45**	-	15,014 PMIDs

(("Macrolides"[mh] OR "Linezolid"[mh]) AND ("Drug Resistance, Microbial"[mh] OR Mutation[mh]))

**Q46**	-	7,165 PMIDs

(("Glycopeptides"[mh] OR "Vancomycin"[mh]) AND ("Drug Resistance, Microbial"[mh] OR genetics[mh]))

**Q47**	-	13,091 PMIDs

(("Genetic Transfer"[mh] OR "Conjugation, Microbial"[mh] OR "Plasmids"[mh]) AND "Drug Resistance, Microbial"[mh])

**Q48**	-	18,913 PMIDs

(("Transposons"[mh] OR "DNA, Bacterial"[mh]) AND ("Drug Resistance, Microbial"[mh] OR resistance[All Fields]))

**Q49**	-	12,384 PMIDs

(("Extended Spectrum beta-Lactamases"[All Fields] OR "AmpC"[All Fields] OR "ESBL"[All Fields]) AND (bacteria[mh] OR genetics[mh]))

**9. Pharmacological substance terms**	(Q50-Q58)

*Substance-indexed queries for anti-infective and antibacterial agents, capturing records indexed pharmacologically rather than by disease or organism descriptor.*

**Q50**	-	29,504 PMIDs

("Anti-Infective Agents"[Substance] AND ("Drug Resistance, Microbial"[mh] OR bacteria[mh]))

**Q51**	-	19,184 PMIDs

("Anti-Infective Agents"[Substance] AND (resistance[All Fields] OR "genetic"[All Fields]))

**Q52**	-	6,897 PMIDs

("Anti-Infective Agents"[Substance] AND (Neisseria meningitidis[mh] OR Streptococcus pneumoniae[mh] OR Haemophilus influenzae[mh] OR Mycobacterium tuberculosis[mh] OR Staphylococcus aureus[mh]))

**Q53**	-	25,647 PMIDs

("Anti-Infective Agents"[Substance] AND ("drug effects"[sh]))

**Q54**	-	10,066 PMIDs

("Anti-Infective Agents"[Substance] AND (susceptibility[All Fields] OR "MIC"[All Fields]))

**Q55**	-	244,826 PMIDs

("Anti-Bacterial Agents"[Substance] AND ("Drug Resistance, Microbial"[mh] OR bacteria[mh]))

**Q56**	-	169,024 PMIDs

("Anti-Bacterial Agents"[Substance] AND (resistance[All Fields] OR "genetic"[All Fields] OR mutation[All Fields]))

**Q57**	-	74,512 PMIDs

("Anti-Bacterial Agents"[Substance] AND (susceptibility[All Fields] OR "MIC"[All Fields] OR "minimum inhibitory concentration"[All Fields]))

**Q58**	-	293,315 PMIDs

("Anti-Bacterial Agents"[Substance] AND ("drug effects"[sh] OR pharmacology[sh]))

**10. Anti-Bacterial Agents (MeSH descriptor)**	(Q59-Q63)

**Q59**	-	178,673 PMIDs

("Anti-Bacterial Agents"[mh] AND ("Drug Resistance, Microbial"[mh] OR resistance[All Fields]))

**Q60**	-	250,643 PMIDs

("Anti-Bacterial Agents"[mh] AND (bacteria[mh] OR bacterial[mh] OR genetics[mh]))

**Q61**	-	30,052 PMIDs

("Anti-Bacterial Agents"[mh] AND (Neisseria meningitidis[mh] OR Streptococcus pneumoniae[mh] OR Haemophilus influenzae[mh] OR Pseudomonas aeruginosa[mh] OR Acinetobacter baumannii[mh]))

**Q62**	-	417,339 PMIDs

("Anti-Bacterial Agents"[mh] AND ("drug effects"[sh] OR pharmacology[sh] OR therapeutic use[sh]))

**Q63**	-	65,257 PMIDs

("Anti-Bacterial Agents"[mh] AND (Mutation[mh] OR genetics[mh] OR gene[All Fields]))

**11. Antibiotic class-specific queries**	(Q64-Q83)

*Per-class coverage of tetracyclines, macrolides, sulfonamides, trimethoprim, chloramphenicol, penicillins, oxazolidinones, lipopeptides, rifamycins, fosfomycin, and polymyxins.*

**Q64**	-	19,416 PMIDs

("Tetracyclines"[mh] OR "Tetracycline Resistance"[mh]) AND ("Drug Resistance, Microbial"[mh] OR bacteria[mh])

**Q65**	-	5,984 PMIDs

(Tetracycline[mh] AND ("Drug Resistance, Microbial"[mh] OR resistance[All Fields]))

**Q66**	-	4,855 PMIDs

(("Doxycycline"[mh] OR "Minocycline"[mh]) AND ("Drug Resistance, Microbial"[mh] OR genetic[All Fields]))

**Q67**	-	26,265 PMIDs

("Macrolides"[mh] AND ("Drug Resistance, Microbial"[mh] OR bacteria[mh] OR genetics[mh]))

**Q68**	-	9,260 PMIDs

(("Erythromycin"[mh] OR "Azithromycin"[mh] OR "Clarithromycin"[mh]) AND ("Drug Resistance, Microbial"[mh] OR resistance[All Fields]))

**Q69**	-	10,705 PMIDs

("Sulfonamides"[mh] OR "Sulfonamide Resistance"[mh]) AND ("Drug Resistance, Microbial"[mh] OR bacteria[mh])

**Q70**	-	3,043 PMIDs

(Sulfamethoxazole[mh] AND ("Drug Resistance, Microbial"[mh] OR resistance[All Fields]))

**Q71**	-	5,073 PMIDs

("Trimethoprim"[mh] OR "Trimethoprim-Sulfamethoxazole"[mh]) AND ("Drug Resistance, Microbial"[mh] OR bacteria[mh])

**Q72**	-	4,443 PMIDs

(("Trimethoprim"[mh] OR "TMP-SMX"[All Fields]) AND (resistance[All Fields] OR genetic[All Fields]))

**Q73**	-	9,430 PMIDs

("Chloramphenicol"[mh] OR "Chloramphenicol Resistance"[mh]) AND ("Drug Resistance, Microbial"[mh] OR bacteria[mh])

**Q74**	-	5,907 PMIDs

(Chloramphenicol[mh] AND (resistance[All Fields] OR genetic[All Fields]))

**Q75**	-	17,168 PMIDs

("Penicillins"[mh] AND ("Drug Resistance, Microbial"[mh] OR "beta-Lactamases"[mh]))

**Q76**	-	11,233 PMIDs

(("Amoxicillin"[mh] OR "Ampicillin"[mh] OR "Penicillin G"[mh]) AND ("Drug Resistance, Microbial"[mh] OR resistance[All Fields]))

**Q77**	-	4,489 PMIDs

(("Oxazolidinones"[mh] OR "Linezolid"[mh]) AND ("Drug Resistance, Microbial"[mh] OR bacteria[mh]))

**Q78**	-	2,872 PMIDs

(Linezolid[mh] AND (resistance[All Fields] OR genetic[All Fields]))

**Q79**	-	12,509 PMIDs

("Lipopeptides"[mh] OR "Daptomycin"[mh]) AND ("Drug Resistance, Microbial"[mh] OR bacteria[mh])

**Q80**	-	11,782 PMIDs

("Rifamycins"[mh] OR "Rifampicin"[mh]) AND ("Drug Resistance, Microbial"[mh] OR bacteria[mh])

**Q81**	-	9,267 PMIDs

(Rifampicin[mh] AND (resistance[All Fields] OR genetic[All Fields] OR mutation[All Fields]))

**Q82**	-	1,780 PMIDs

("Fosfomycin"[mh]) AND ("Drug Resistance, Microbial"[mh] OR bacteria[mh] OR resistance[All Fields])

**Q83**	-	7,885 PMIDs

(("Polymyxins"[mh] OR "Colistin"[mh] OR "Polymyxin B"[mh]) AND ("Drug Resistance, Microbial"[mh] OR bacteria[mh]))

**12. Regulatory mechanisms, enzymatic inactivation, and nitroimidazoles**	(Q84-Q87)

**Q84**	-	5,753 PMIDs

(("two-component" OR "regulatory system" OR "sensory histidine kinase" OR "response regulator" OR "quorum sensing")[Title/Abstract] AND (resistance[Title/Abstract] OR "drug effect"[Title/Abstract]))

**Q85**	-	78,192 PMIDs

(("signal transduction" OR "cell envelope" OR "outer membrane" OR "lipopolysaccharide" OR "peptidoglycan")[Title/Abstract] AND (mutation[Title/Abstract] OR resistance[Title/Abstract]))

**Q86**	-	1,887 PMIDs

(("enzymatic inactivation" OR "chemical modification" OR "antibiotic inactivation")[Title/Abstract] AND (resistance[Title/Abstract] OR gene[Title/Abstract]))

**Q87**	-	6,243 PMIDs

(("Nitroimidazoles"[mh] OR "Metronidazole"[mh]) AND ("Drug Resistance, Microbial"[mh] OR bacteria[mh]))

**Section S2. Corpus assembly - Stage 2: metadata pre-filtering**

Before language-model relevance screening, the candidate corpus assembled in Stage 1 was reduced to a bacterial, AMR-plausible subset by a four-signal classifier operating over structured metadata. This section documents the two metadata sources, the four pass signals, the post-filter organism and language cleanup, and the complete controlled-vocabulary term lists used by each signal.

**S2.1 Metadata sources**

Two metadata sources were assembled for every candidate PMID. Entity annotations were drawn from the PubTator3 bulk annotation files (gene, chemical, species, mutation, and relation), which were streamed and indexed into a per-PMID table of boolean flags recording whether a gene, chemical, species, or mutation was mentioned, together with a flag marking the presence of a high-value relation pair. High-value relation pairs were defined as gene-chemical, variant-chemical, gene-disease, and variant-disease co-annotations, these being the relation types most indicative of a resistance-mechanism report. Bibliographic metadata (abstract, MeSH descriptors and their topical subheadings, indexed substances, publication types, and language) were retrieved through the Entrez API and stored in a local database keyed on PMID.

**S2.2 Pass signals**

A paper was retained if any one of the following four signals fired. The signals are deliberately redundant: a record indexed under any single scheme is kept, so that incomplete annotation under one scheme does not exclude a genuinely relevant paper. This is consistent with the pipeline's high-recall posture at all stages preceding structured extraction.

**Signal 1 - PubTator3 co-mention.** A gene or mutation is co-mentioned with a chemical in the same record (has_gene OR has_mutation) AND has_chemical. Records additionally carrying a high-value relation pair, or a mutation mention, are flagged as higher-priority within the pass set.

**Signal 2 - MeSH descriptor co-occurrence.** An AMR-mechanism MeSH descriptor (S2.3) co-occurs with an antibiotic or resistance-testing descriptor (S2.4).

**Signal 3 - MeSH subheading.** A genetic-characterisation MeSH subheading (S2.5) co-occurs with an AMR-associated indexed substance (S2.6) or an AMR-mechanism descriptor.

**Signal 4 - species with mechanism and drug.** A PubTator3 species mention co-occurs with both an AMR-mechanism descriptor and a drug or resistance descriptor.

**S2.3 Post-filter cleanup**

Records passing any signal were then cleaned in two steps. Non-English records were removed on the basis of the Entrez language field. Records whose MeSH terms indicated a non-bacterial subject (S2.7) were removed only where no confirming bacterial term (S2.8) was also present, so that papers of mixed subject (for example a comparative study naming both bacterial and fungal resistance) were retained.

**S2.4 Term lists**

The controlled-vocabulary sets used by the signals are reproduced below in full, exactly as applied. MeSH matching was performed on descriptor main headings; subheading matching was case-insensitive.

**AMR-mechanism MeSH descriptors (Signals 2, 3, 4)**

**Descriptors (48).** ATP-Binding Cassette Transporters, Acetyltransferases, Bacterial Outer Membrane Proteins, Bacterial Persistence, Bacterial Proteins, Bacterial Transcription, Bacteriophages, Biofilms, Carbapenemases, Conjugation, Microbial, Cross Resistance, DNA Gyrase, DNA Topoisomerase IV, DNA Transposable Elements, Drug Resistance, Bacterial, Drug Resistance, Microbial, Drug Resistance, Multiple, Bacterial, Frameshift Mutation, Gene Cassettes, Gene Dosage, Gene Expression Regulation, Bacterial, Genes, Bacterial, Genes, Essential, Genetic Transfer, Horizontal, Integrons, Membrane Transport Proteins, Metallo-beta-Lactamases, Methicillin Resistance, Methyltransferases, Microbial Drug Resistance, Microbial Viability, Multidrug Resistance-Associated Proteins, Multigene Family, Mutation, Mutation, Missense, Operons, Penicillin Resistance, Penicillin-Binding Proteins, Phosphotransferases, Plasmids, Point Mutation, Porins, Prophages, Regulon, Ribosomal Proteins, Transformation, Bacterial, Transposons, beta-Lactamases

**Drug and resistance MeSH descriptors (Signals 2, 4)**

**Descriptors (69).** Amikacin, Aminoglycosides, Amoxicillin, Ampicillin, Anti-Bacterial Agents, Anti-Infective Agents, Antibiotics, Azithromycin, Carbapenems, Cefepime, Cefotaxime, Ceftazidime, Ceftriaxone, Cephalosporins, Chloramphenicol, Ciprofloxacin, Clarithromycin, Clindamycin, Colistin, Daptomycin, Disk Diffusion Antimicrobial Tests, Doripenem, Doxycycline, Drug Resistance, Ertapenem, Erythromycin, Fluoroquinolones, Fosfomycin, Fusidic Acid, Gentamicin, Glycopeptides, Imipenem, Kanamycin, Lactams, Levofloxacin, Linezolid, Macrolides, Meropenem, Metronidazole, Microbial Sensitivity Tests, Minimum Inhibitory Concentration, Minocycline, Monobactams, Moxifloxacin, Mupirocin, Nitrofurantoin, Nitroimidazoles, Oxazolidinones, Penicillins, Piperacillin, Polymyxin B, Polymyxins, Quinolones, Rifampicin, Rifampin, Rifamycins, Streptomycin, Sulfamethoxazole, Sulfonamides, Tedizolid, Teicoplanin, Tetracycline, Tetracyclines, Tigecycline, Tobramycin, Trimethoprim, Vancomycin, beta-Lactamase Inhibitors, beta-Lactams

**Genetic-characterisation MeSH subheadings (Signal 3)**

**Subheadings (12).** biosynthesis, classification, drug effects, enzymology, genetics, isolation, isolation & purification, metabolism, microbiology, pathogenicity, pharmacology, physiology

**AMR-associated indexed substances (Signal 3)**

**Substances (19).** AmpC beta-Lactamases, Anti-Bacterial Agents, Anti-Infective Agents, Bacterial Proteins, DNA, Bacterial, Efflux Pumps, Extended-Spectrum beta-Lactamases, Integrons, Membrane Proteins, Metallo-beta-Lactamases, Penicillin-Binding Proteins, Plasmids, Porins, RNA, Bacterial, RNA, Ribosomal, 16S, RNA, Ribosomal, 23S, beta-Lactamases, beta-Lactams, carbapenemase

**Non-bacterial subject terms (cleanup, removal)**

**Terms (42).** Antifungal Agents, Antiparasitic Agents, Antiviral Agents, Aspergillus, Aspergillus fumigatus, Candida, Candida albicans, Candida auris, Candida glabrata, Coronavirus, Cryptococcus, Cryptococcus neoformans, Cryptosporidium, Cytomegalovirus, Dermatophytes, Drug Resistance, Fungal, Drug Resistance, Parasites, Drug Resistance, Viral, Fungi, Fusarium, Giardia, HIV, HIV-1, HIV-2, Helminths, Hepatitis B virus, Hepatitis C virus, Herpesvirus, Influenza A virus, Influenza B virus, Leishmania, Malaria, Plasmodium, Plasmodium falciparum, Plasmodium vivax, Pneumocystis, SARS-CoV-2, Saccharomyces, Toxoplasma, Trichophyton, Trypanosoma, Viruses

**Bacterial-confirmation terms (cleanup, override)**

**Terms (35).** Acinetobacter, Acinetobacter baumannii, Anti-Bacterial Agents, Bacteria, Bacterial Proteins, Bacteroidetes, Burkholderia, Campylobacter, Clostridioides difficile, Clostridium, Drug Resistance, Bacterial, Drug Resistance, Multiple, Bacterial, Enterobacteriaceae, Enterococcus, Escherichia coli, Gene Expression Regulation, Bacterial, Genes, Bacterial, Gram-Negative Bacteria, Gram-Positive Bacteria, Haemophilus, Helicobacter pylori, Klebsiella, Klebsiella pneumoniae, Listeria, Mycobacterium, Mycobacterium tuberculosis, Neisseria, Pseudomonas, Pseudomonas aeruginosa, Salmonella, Staphylococcus, Staphylococcus aureus, Stenotrophomonas, Streptococcus, beta-Lactamases

**Section S3. Stage 3: relevance screening**

The corpus surviving metadata pre-filtering was screened for AMR relevance by a BioMistral-7B language model under few-shot prompting. This section documents the record fields presented to the classifier, the system instruction and few-shot exemples, the decoding and decision rule, and the substance keyword list used to append an advisory line to the prompt. Together these allow the screening step to be reproduced exactly.

**S3.1 Input record and decision rule**

Screening read the structured records held in the local database populated during earlier stages. Records without an abstract were excluded at query time and did not enter screening. For each remaining record the classifier was presented with the fields below, assembled into a single text block; fields absent for a given record were omitted from its block.

```
Title: <article title>
Journal: <journal>
Abstract: <abstract>
MeSH Terms: <mesh terms>        [line omitted if field empty]
Substances: <indexed substances> [line omitted if field empty]
Keywords: <author keywords>      [line omitted if field empty]
Extracted Entities - Has Gene: <bool>, Has Chemical: <bool>, Has Mutation: <bool>
Note: Substances suggest possible resistance mechanism.   [conditional; see S3.4]
```

The block was preceded by the system instruction and five worked examples (S3.2-S3.3) and closed with the question “Is this paper about antimicrobial resistance? Answer:”. Inference used greedy decoding (do_sample = False) in half precision, generating at most ten new tokens. The generated continuation was truncated at the first newline, any “Answer:” prefix removed, and the record classified as AMR-relevant if the remaining text contained “YES” (case-insensitive). Prompts were left-padded and processed in sub-batches; the prompt set was divided into contiguous blocks, one per available GPU, each processed by an independent worker holding its own model replica.

**S3.2 System instruction**

The system instruction, prepended to every prompt, was:

You are an expert microbiologist reviewing papers for antimicrobial resistance (AMR). This includes: antibiotic resistance genes and mutations, drug target genes (DHPS, DHFR, GyrA, PBPs), gene cassettes, integrons, transposons, mobile genetic elements, efflux pumps, beta-lactamases, and susceptibility studies. A paper is relevant even if it does not use the word 'resistance' - cloning or characterizing an AMR-related gene, transposon, or its mutations counts as YES. Papers about resistance-gene-carrying transposons are YES, even if written in highly technical molecular biology language (e.g., cointegrate formation, IS elements). Answer strictly YES or NO.

**S3.3 Few-shot exemples**

Five worked examples were prepended to each query record, each rendered in the same field format as the query and each labelled “YES”. All five are positive instances; no negative example was supplied. This tunes the classifier toward high recall, consistent with the pipeline's precision-over-recall allocation, in which false positives are removed during structured extraction (Section S5) whereas false negatives are unrecoverable. The example for PMID 11557503 additionally carried an inline note stating that gene cassettes and integrons indicate AMR even where no gene entities were extracted; the other four carried no such note.

**Table S3. Few-shot example PMIDs.**

| **PMID** | **Role** |
| --- | --- |
| 10027979 | Positive example (label: YES) |
| 10049269 | Positive example (label: YES) |
| 11557503 | Positive example (label: YES); inline note on gene cassettes / integrons |
| 1522070 | Positive example (label: YES) |
| 12726767 | Positive example (label: YES) |

**S3.4 Substance advisory terms**

Where a record's indexed-substance field contained any of the terms below (case-insensitive substring match), a single advisory line - “Note: Substances suggest possible resistance mechanism.” - was appended to that record's prompt block. The advisory was informational and did not by itself determine the classification.

**Terms (49).** resistance, resistant, susceptibility, susceptible, MIC, lactamase, carbapenemase, acetyltransferase, phosphotransferase, nucleotidyltransferase, esterase, efflux, pump, transporter, porin, permeability, methyltransferase, demethylase, integron, cassette, plasmid, transposon, mobilizable, conjugative, insertion sequence, operon, allele, variant, genotype, SNP, mutation, missense, point mutation, aminoglycoside, tetracycline, methicillin, vancomycin, colistin, polymyxin, carbapenem, cephalosporin, fluoroquinolone, macrolide, sulfonamide, oxazolidinone, glycopeptide, penicillin, fosfomycin, chloramphenicol

**Section S4. Full-text retrieval**

Papers screened as AMR-relevant (Section S3) were passed to a custom full-text retrieval pipeline for downstream extraction. The pipeline queries only open, publicly documented APIs and web endpoints - no institutional proxy, no third-party scraping service, and no browser automation in the production path. The pipeline requires no paid subscription, though optional publisher API keys (Elsevier, Wiley, Springer). It is implemented in Python 3, and runs in three sequential stages: open-access retrieval, publisher- and API-mediated retrieval for records the open-access stage could not resolve, and a merge/deduplication/validation stage. This section documents the retrieval routes, the full-text validation that distinguishes genuine article bodies from preview and placeholder pages, and the resulting corpus composition.

**S4.1 Retrieval cascade**

Each PMID is first enriched with metadata (title, journal, date, DOI) via NCBI E-utilities esummary, and mapped to a PMCID via the PMC ID Converter. Two ordered cascades are then attempted, stopping at the first route that returns a validated article body (S4.2).

Stage 1 - open-access routes, in strict priority order:

**1.** PMC Open Access bulk archive - the publisher's own structured JATS XML, the highest-fidelity source.

**2.** PMC OA subset via the oa.fcgi web service, fetched as BioC JSON and flattened into labelled sections.

**3.** Europe PMC full-text XML, which often exposes open-access articles absent from NCBI's own OA subset.

**4.** NCBI PMC EFetch XML, accepted only if the returned body exceeds 200 characters (publishers return abstract-only stubs through this endpoint).

**5.** PMC rendered article HTML, converted to text and validated to reject PDF-viewer shell pages.

**6.** DOI-based open-access discovery: the DOI is resolved via Crossref and enriched with Unpaywall, and Europe PMC OA locations, and each candidate landing page or PDF is fetched and validated, with journal-specific URL rewriting applied first.

PMIDs that exhaust all six routes, and those resolving only to abstract/metadata, are queued for Stage 2.

Stage 2 - publisher- and API-mediated routes, run only on the unresolved set. Stage 2 retries the highest-value PMC and Europe PMC routes (guarding against transient failure) and then works through a publisher-aware cascade with explicit per-publisher URL construction and host allow-listing for Wiley, Oxford University Press, ASM, SAGE, ACS, Springer/Nature, PLOS, J-STAGE, RSC, Eurosurveillance, Cambridge, and Microbiology Society journals. Publisher-specific handlers include a dedicated cookie-based SAGE session with a policy-matched rate limiter; the Springer Nature OA API and Elsevier Article API (both key-gated); ASM and Wiley text-and-data-mining (TDM) endpoints. PDF-only retrievals (Wiley TDM, generic publisher PDF) are archived but explicitly flagged as non-body content so they cannot be counted as full text. PubMed BioC serves as the terminal fallback, guaranteeing at least title, abstract, and metadata for every PMID with a DOI. Every attempted backend and every backend-level error is recorded per PMID, and abstract-only outcomes are bucketed by reason (e.g. publisher gate, Elsevier entitlement, legacy Wiley) to guide future access-expansion work.

**S4.2 Full-text validation**

Because many routes return syntactically successful but practically useless content (preview pages, PDF-viewer shells, metadata-only XML, or PDF placeholders) every retrieval passes a second, stricter validation independent of retrieval success. The validation is source-aware. For structured XML, a record is accepted only if it contains at least one genuine body section (excluding abstract, keywords, funding, acknowledgements, and references by an explicit name-list) and the body exceeds a word-count threshold (default 500 words). Elsevier XML must additionally contain real section elements, rejecting entitlement-blocked metadata responses. HTML-derived sources are screened by heuristic detectors that reject PMC PDF-viewer shells, publisher preview/loading shells, and journal-overview pages; where recognised section headings are present the inter-heading body length is used, and where none are found a much higher word bar (≥1,800 words) is required before the page is trusted. Wiley TDM and Europe PMC PDF placeholders are always rejected, since only a PDF was archived and the pipeline performs no PDF-to-text extraction. This logic runs both inline during retrieval and as an independent post-hoc pass over the final corpus, so corpus composition can be re-derived or re-thresholded from the downloaded files without re-issuing any network request.

All retrieved content is normalised to a single plain-text schema, a metadata header followed by “## SECTION” headings and body text, so downstream extraction sees one consistent format regardless of source. When both an open-access and an API-stage file exist for a PMID, the higher-priority source wins under a fixed ranking (PMC OA bulk XML > Europe PMC XML > PMC EFetch XML > PMC OA BioC > Elsevier XML > Elsevier text > OA HTML > publisher HTML > publisher PDF > PubMed BioC). Validated articles are written to the full-text corpus; everything else is retained as abstract-only.

**S4.3 Rate limiting and reproducibility**

All outbound requests pass through a single rate limiter enforcing NCBI guidance (3 req/s without an API key, 10 req/s with one) plus a policy-matched SAGE override, with exponential-backoff retry on HTTP 429 and a descriptive User-Agent and contact email on every NCBI call. Publisher credentials (NCBI, Elsevier, Wiley TDM, Springer) are loaded from a git-ignored environment file; none is required for the core PMC/Europe PMC/Crossref/Unpaywall routes.

**Section S5. Stage 5: structured extraction**

This section documents the two-pass extraction stage: the model and inference parameters, the output schema, the deterministic filters applied before and after auditing, and both prompts reproduced in full. Together these allow the extraction of genes and mutations from full text to be reproduced and audited.

**S5.1 Model and inference parameters**

Extraction used Qwen3-30B-A3B, a mixture-of-experts model, served with vLLM in bfloat16 precision on NVIDIA GH200 nodes at the TACC Vista cluster. Both passes decoded greedily with reasoning mode disabled. All parameters were fixed before inference and applied unchanged to every document.

| **Parameter** | **Pass 1 - extraction** | **Pass 2 - audit** |
| --- | --- | --- |
| Model | Qwen3-30B-A3B | Qwen3-30B-A3B |
| Serving framework | vLLM | vLLM |
| Precision | bfloat16 | bfloat16 |
| Context window | 32,768 tokens | 32,768 tokens |
| Tensor parallel size | 1 | 1 |
| GPU memory utilisation | 0.92 | 0.92 |
| Temperature | 0.0 | 0.0 |
| top-p | 1.0 | 1.0 |
| Repetition penalty | 1.1 | 1.05 |
| Max output tokens | 12,000 whole paper; 8,000 per chunk | 4,096 per entity |
| Reasoning mode | disabled | disabled |

Papers of 66,360 characters or fewer were extracted in a single pass. Longer papers were split into 55,000-character chunks with 1,000-character overlap, extracted independently, and merged, with mutation entries deduplicated across chunks. Audit context was capped at 80,000 characters; where a paper exceeded this, the auditor was given the first 8,000 characters plus 4,000-character windows centred on each mention of the entity under review, overlapping windows merged. Papers were processed one at a time, each producing an individual JSON file so that interrupted runs resume without reprocessing.

**S5.2 Output schema**

Schema enforcement is strict: fields absent from the model output receive the schema default, null lists become empty lists, and any key not defined in the schema is dropped. Gene entries never carry a mutations_found field - where the model emitted one, its contents were relocated to the mutations collection before enforcement rather than discarded.

Paper-level record: pmid, relevant, paper_title, publication_year, paper_type (single_gene, multi_gene_resistome, review, or computational_with_validation), genes, mutations, sequence_accessions, key_findings, methodology, geographic_location, sample_size.

**Gene entry**

allele, encodes, mechanism, confers_resistance_to, resistance_mechanism_class, organisms_tested_in, role_in_paper, validation_method, evidence_level, key_substitutions, genetic_context, source, evidence_note.

**Mutation target and nested mutation entry**

Target: encodes, mutations_found, wild_type_susceptible, notes, source, evidence_note. Each entry in mutations_found: notation, amino_acid_position, nucleotide_position, codon_change, nucleotide_change, protein_change, position_in_molecule, confers_resistance_to, organisms_observed_in, effect_on_function, mutation_type, validated_by, origin, evidence_level.

The model intermittently emitted alternative field names (e.g. “substitution” for notation, “position” for amino_acid_position, “domain” for position_in_molecule). These were folded into the canonical fields before enforcement. Folding is context-aware: where the position or type indicated a non-coding region (promoter, Pribnow box, -10/-35 element, RBS, UTR, intergenic), protein-level aliases were redirected to nucleotide-level fields, preventing a promoter variant from being recorded as an amino-acid substitution.

**S5.3 Genes barred from the gene catalogue**

The following confer resistance only when mutated or inactivated. They were routed to the mutations catalogue and blocked from the genes catalogue both by prompt rule and by a deterministic denylist applied after extraction.

| **Category** | **Genes** |
| --- | --- |
| Two-component systems | pmrA/B, phoP/Q, parR/S, colR/S, cprR/S, rppA/B, qseB/C, basR/S, envZ/ompR |
| Negative regulators | mgrB, mgrR, micA |
| LPS biosynthesis and lipid A remodelling | lpxA-T, arnB-T, pmrC, pmrE, eptA-C, cptA, ugd, waaL, waaP, rfbA, lpsB, galU |
| Acylation / outer membrane | msbB, pagP, pagL |
| LPS transport | lptA-G |
| Sigma factors | rpoN, rpoE, rpoS, rpoH, rpoD |
| Efflux regulators | mexR, nalC, nalD, mexZ, mexT, acrR, marR/marA, soxR/soxS, ramR/ramA |
| Hopanoid biosynthesis | isph, hpnJ |
| rRNA loci | 16S, 23S, 12S and variant spellings |

Operon-name prefixes (arnBCAD, pmrCAB, lpxABC, phoPQ, pmrAB, parRS, colRS) and locus-tag patterns (PA####, PA14_####, ORF####, locus_####) were excluded by the same mechanism. Chromosomal core genes whose mutation confers resistance (gyrA, parC, rpoB, rpsL, folA, ompK35/36 and the penicillin-binding proteins) are excluded by prompt rule. No positive whitelist of gene names was applied: restricting the catalogue to an enumerated list would exclude newly named determinants from discovery papers, which are among the entries of greatest interest to curators. Precision is instead enforced by the denylist, the prompt validation gate, the audit pass, and a final check that each gene name appears literally in the source text.

**S5.4 Drug-class filtering**

Antibiotic assignments were filtered against the gene-family-to-drug-class map below. Assignments outside a family's permitted classes were removed; genes outside these nineteen families passed unfiltered. Matching was case-insensitive and substring-based, so truncated stems match their full drug names.

| **Family** | **Permitted antibiotics and classes** |
| --- | --- |
| bla | penicillins, cephalosporins, carbapenems, monobactams, and named β-lactams (amoxicillin, ampicillin, piperacillin, cefotaxime, ceftazidime, imipenem, meropenem, aztreonam) |
| aac / aph / aad / ant | aminoglycosides (gentamicin, tobramycin, amikacin, kanamycin, streptomycin, spectinomycin) |
| tet | tetracycline, doxycycline, minocycline, tigecycline |
| qnr | quinolones, fluoroquinolones |
| erm / mef / mph | macrolides (± lincosamides, streptogramin for erm) |
| van | vancomycin, teicoplanin, glycopeptide |
| cfr | phenicols, oxazolidinones, lincosamides, pleuromutilins, streptogramin |
| optrA / poxtA | oxazolidinones, phenicols |
| cat | chloramphenicol, phenicol |
| sul | sulfonamides |
| dfr | trimethoprim |
| mcr | colistin, polymyxin |
| fos | fosfomycin |

Multi-class families were preserved in full (cfr across the MOLPS classes; erm across the MLS group). Vague claims (“multidrug”, “MDR”, “multiple antibiotics”, “broad spectrum”, and unqualified “beta-lactams”) were removed wherever they appeared. The full prefix-to-antibiotic listing is in the code repository.

**S5.5 β-lactamase name normalisation**

Bare enzyme names (ADC-56, OXA-23, CTX-M-15) were resolved to bla-prefixed gene names by pattern rather than enumerated list, so the rule extends to families not yet described. A name of two to six letters, optionally with an -M infix, followed by a hyphen and a number, was treated as a β-lactamase allele unless its leading letters matched one of 44 non-β-lactamase AMR gene prefixes (aac, aph, tet, qnr, van, mcr, gyr, par, rpo, pbp and others). Thus ADC-56 → blaADC-56 and CTX-M-15 → blaCTX-M-15, while aac(6')-Ib (parentheses), tet-1 (excluded stem) and gyrA (no allele number) are left unchanged. A related step lifts β-lactamase alleles misfiled as mutation targets into the gene catalogue where explicit coding substitutions are present; promoter and regulatory variants are left in the mutation catalogue.

**S5.6 Review-paper handling**

Papers were classified as reviews when two or more of sixteen signal phrases (“this review”, “we review”, “minireview”, “an overview of”, “current understanding”, and similar) appeared in the first 3,000 characters. Entries from review papers were marked source “bibliography” and evidence_level “inferred”, with codon changes, nucleotide changes and nucleotide positions forced to null, since such papers report other groups' findings rather than original data.

**S5.7 Deterministic post-processing sequence**

Filters were applied in fixed order, before and after the audit pass.

| **Order** | **Filter** | **Action** |
| --- | --- | --- |
| 1 | β-lactamase reconciliation | Lift misfiled alleles from mutations to genes where coding substitutions are explicit |
| 2 | Gene-key normalisation | Resolve bare enzyme names to bla-prefixed form; merge duplicates |
| 3 | Mutation promotion | Relocate gene-level mutations_found arrays to the mutations collection |
| 4 | Denylist | Remove regulators, biosynthesis genes, operon names, locus tags from genes |
| 5 | Schema enforcement | Apply canonical shapes; drop non-schema keys; normalise null lists |
| 6 | Mutation cleanup | Remove blocked targets, empty entries, targets with no usable mutations |
| 7 | Drug-class filter | Remove off-class antibiotics and vague multidrug claims |
| 8 | Atomic audit | Per-entity verification against source text (Pass 2) |
| 9 | Filters 1-7 reapplied | Post-audit normalisation of audited output |
| 10 | Literal-name check | Remove genes whose name does not appear in the source text |
| 11 | Final sweep | Remove genes with neither a specific antibiotic nor a mechanism class |

The literal-name check was skipped for computational_with_validation papers, where genes are legitimately named by internal family codes that may appear only in figures. The final sweep required both conditions to fail: a gene with a known mechanism class was retained even where no specific antibiotic survived auditing.

**S5.8 Extraction prompt (Pass 1)**

Reproduced as executed. {chunk_note} and {paper_text} are runtime placeholders. The prompt was locked before inference and applied unchanged to every document.

```
You are an antimicrobial resistance (AMR) extraction system. Read the paper text below and output a JSON object describing the AMR genes and mutations characterized in THIS paper.

OUTPUT: a single JSON object only. Start with { immediately. No prose, no markdown.
{chunk_note}
================================================================================
SECTION 1 - SCOPE OF THE "genes" SECTION
================================================================================
The "genes" section holds acquired or intrinsic AMR genes whose wild-type
product directly confers resistance - i.e., the gene works on its own, no
mutation required. Examples of families that qualify:
  - Beta-lactamases (bla-prefixed enzymes)
  - Aminoglycoside-modifying enzymes (aac, aph, aad, ant)
  - Tetracycline / quinolone / macrolide / lincosamide / streptogramin
    resistance genes (tet, qnr, erm, mef, mph, lnu, lsa, vat, vgb, ...)
  - Phenicol / oxazolidinone resistance (cat, cfr, optrA, poxtA, ...)
  - Glycopeptide cassettes (vanA-G), polymyxin (mcr), fosfomycin (fos)
  - Sulfonamide / trimethoprim (sul, dfr)
  - Acquired efflux pumps reported as full operons (mexAB-OprM, acrAB-TolC,
    oqxAB, ...)
  - Novel AMR genes characterized for the first time in the paper, named by
    the authors (e.g. a paper describing a new bacitracin amidohydrolase
    might name it bahA). Include these when the paper experimentally
    demonstrates the resistance phenotype.

NAMING CONVENTION:
- Beta-lactamases: use the canonical "bla<FAMILY>-<N>" form as the JSON key,
  even when the paper writes the enzyme bare. The bare form names the enzyme;
  the gene name carries the "bla" prefix.
- Other AMR genes: use the name as written, preserving punctuation
  (e.g. "aac(6')-Ib", "tet(M)", "qnrS1").

What does NOT belong in "genes" (these go in "mutations" instead):
- Regulators whose mutation/loss is the resistance mechanism (TCS like
  pmrA/B, phoP/Q, parR/S, mgrB; efflux regulators like mexR, acrR; sigma
  factors; etc.)
- Biosynthesis / cell-envelope genes (lpxA-T, arnB-T, eptA-C, waaL, pagP,
  cell-wall remodeling enzymes)
- Chromosomal core genes whose mutation creates resistance (gyrA, parC,
  rpoB, rpsL, folA, ompK35/36, PBPs, rRNAs)
- Locus tags (PA####, ORF####, locus_####)

Special allowance - when genes are named with internal family codes such
as "C1264", "E449", use as gene name the overall gene family and the specific name as
allele. When the paper validates them experimentally, extract
each validated gene using its internal identifier as the key.

================================================================================
SECTION 2 - THE RESULTS-ONLY RULE (single most important filter)
================================================================================
Extract ONLY genes and mutations that represent THIS paper's OWN findings.
A gene belongs in the output if and only if it appears in:
  ✓ The Results section with experimental data generated in this study
  ✓ A table of THIS paper's own results (MIC, gene characterization, mutations)
  ✓ The Methods section as a gene cloned or assayed in this study
  ✓ The abstract as a finding of this study

EXCLUDE genes that appear only in:
  ✗ Introduction as background or context ("X was previously reported...")
  ✗ Discussion as comparison to other work ("similar to what Y et al. found...")
  ✗ Reference list citations
  ✗ Any sentence containing: "previously", "reported by", "described by",
    "identified by", "according to", "et al.", "was shown to", "has been"
    when referring to other groups' work

THE TEST: "Did the authors of THIS paper generate new data about this gene?"
  YES → include    NO → exclude

EXCEPTION - allele-characterization papers:
For papers that name new alleles AND deposit sequences AND perform functional tests
(hydrolysis, MIC on transformant, or susceptibility testing of the native strain),
extract ONLY the alleles that were functionally tested, NOT the full WGS-discovered list.
WGS-only discovery without any functional data does NOT qualify.

================================================================================
SECTION 3 - VALIDATION GATE (the second filter)
================================================================================
DIRECTION OF RESISTANCE CHECK:
AMR genes and mutations confer RESISTANCE - they reduce susceptibility of a
microorganism to a drug. If the experimental finding is the OPPOSITE -
increased susceptibility, hypersusceptibility, sensitization, or enhanced
drug toxicity - the entry does NOT belong in genes or mutations.
This applies regardless of how thoroughly the phenotype is characterized.

A gene passes into the "genes" section only if BOTH conditions are met:
  (a) It was experimentally validated in THIS paper AND the validation demonstrated
    an ANTIMICROBIAL RESISTANCE PHENOTYPE - meaning a change in MIC, zone of
    inhibition, growth in the presence of an antibiotic, or enzyme activity against
    an antibiotic substrate. The following do NOT qualify as AMR validation:
      - Macrophage survival or intracellular survival assays
      - Apoptosis induction assays
      - Animal infection models (LD50, mucin model, etc.)
      - Invasion or adherence assays
      - Virulence phenotypes of any kind
      - Complementation of a virulence defect
    A gene characterized only for its role in pathogenesis, metabolism, or host
    interaction MUST NOT appear in "genes" even if extensively validated.
  (b) The wild-type gene confers resistance WITHOUT requiring a mutation
      (i.e., it is an acquired or intrinsic resistance gene, not a regulator
      whose loss-of-function alters susceptibility)

If (b) fails - i.e., the gene confers resistance only when mutated or inactivated -
the entry belongs in the MUTATIONS section, not the genes section.

================================================================================
SECTION 3b - REGULATORY / LOSS-OF-FUNCTION / BIOSYNTHESIS GENES
================================================================================
These genes ALWAYS go in the MUTATIONS section, never the genes section, because
the wild-type does not confer resistance - only mutations or inactivation do:

  Two-component systems: pmrA, pmrB, phoP, phoQ, parR, parS, colR, colS,
    cprR, cprS, rppA, rppB, qseB, qseC, basR, basS, envZ, ompR
  Negative regulators: mgrB, mgrR, micA
  LPS biosynthesis: lpxA, lpxC, lpxD, lpxE, lpxF, lpxH, lpxK, lpxL, lpxM,
    lpxO, lpxP, lpxT, arnB, arnC, arnD, arnE, arnF, arnT, pmrC, pmrE,
    eptA, eptB, eptC, cptA, ugd, waaL, waaP, rfbA, lpsB, galU
  Acylation: msbB, pagP, pagL
  LPS transport: lptA-G
  Sigma factors: rpoN, rpoE, rpoS, rpoH, rpoD
  Efflux regulators: mexR, nalC, nalD, mexZ, mexT, acrR, marR/marA, soxR/soxS,
    ramR/ramA
  Hopanoid: isph, hpnJ
  Locus tags: PA####, PA14_####, ORF####, locus_#### - never extract as genes

Also: language patterns "mutations in X", "inactivation of X", "loss-of-function
in X", "disruption of X", "ΔX confers" → mutations section regardless of family.

================================================================================
SECTION 4 - DRUG CLASS RULES (assignment must match family)
================================================================================
HARD EXCLUSIONS (these are common hallucinations):
  - bla genes do NOT confer fluoroquinolone, colistin, vancomycin, or
    aminoglycoside resistance
  - aac/aph/aad genes do NOT confer beta-lactam, fluoroquinolone, or
    colistin resistance
  - qnr genes do NOT confer beta-lactam or vancomycin resistance
  - tet genes do NOT confer beta-lactam or vancomycin resistance
  - mcr genes confer ONLY colistin / polymyxin resistance

Legitimate multi-class genes (preserve all valid classes):
  - cfr: MLS + oxazolidinones + phenicols + pleuromutilins (MOLPS)
  - optrA, poxtA: oxazolidinones + phenicols
  - erm: macrolides + lincosamides + streptogramin B (MLS)
  - broad efflux pumps (mexAB-OprM, acrAB-TolC): multiple classes ARE valid

Prefer to use the specific antibiotic name(s) compared to vague terms alone ("multidrug", "multiple antibiotics",
"beta-lactams").

================================================================================
SECTION 5 - TABLE EXTRACTION (MANDATORY)
================================================================================
Tables are the PRIMARY source of structured AMR data. Scan EVERY table before
writing JSON. Failure to extract from tables is the most common error.

FOR EVERY TABLE:
  1. Read every row AND every column header
  2. Identify which columns contain gene/allele names, mutation notations,
     MIC values, organism names, or antibiotic names
  3. Extract EVERY named gene or mutation that passes the validation gate
  4. Do not stop at the first family - if a table has OXA, ADC, NDM, KPC
     columns, all families must be extracted

MUTATION TABLES specifically:
  - Every row → a separate mutation entry
  - Column headers map to fields: position → amino_acid_position,
    substitution → notation + protein_change, MIC → confers_resistance_to
  - Different genes in different columns → separate entries in mutations section

GENE CHARACTERIZATION TABLES:
  - Cloned genes with MIC values → extract each gene with specific MICs as
    confers_resistance_to
  - Enzyme kinetics tables → extract Km/kcat if present

DO NOT extract from:
  - Whole-isolate MIC tables (MIC for entire clinical isolate, not specific
    cloned gene) → do not use for confers_resistance_to
  - Epidemiological tables with isolate metadata only
  - Reference tables citing other studies

================================================================================
SECTION 6 - ANTI-HALLUCINATION & NON-CODING MUTATIONS
================================================================================
- Do NOT invent codon changes, nucleotide changes, positions, or organisms.
- If the paper does not state a field explicitly, set it to null (do not guess).
- Do NOT copy mutation notations verbatim across different genes.
- Do NOT generate gene entries that share identical (encodes, mechanism,
  confers_resistance_to, organisms, validation_method) tuples - this is the
  copy-paste loop signature.
- If a gene name appears only in the bibliography or a reference list, exclude it.

For mutations in NON-CODING regions (promoter, RBS, UTR, intergenic - anywhere
outside the protein-coding sequence):
  - protein_change = null, amino_acid_position = null
  - Use nucleotide_change and nucleotide_position
  - position_in_molecule describes the regulatory region
  - The associated gene appears in BOTH "genes" (the wild-type enzyme is the
    AMR gene) and "mutations" (the regulatory change is the mechanism).

================================================================================
SECTION 7 - COMPUTATIONAL + EXPERIMENTAL VALIDATION PAPERS
================================================================================
For papers that computationally discover new gene families AND experimentally
validate representative genes (cloning + MIC or disk diffusion):
  - Set paper_type = "computational_with_validation"
  - Extract EACH experimentally validated gene as its own entry in "genes" using as gene name the overall gene family in which it belongs
  - confers_resistance_to = the actual phenotype observed in validation
  - validation_method = brief description (e.g. "cloning in E. coli + disk diffusion")
  - evidence_level = "computational_with_wet_lab_validation"
  - role_in_paper = "experimentally_characterized"
  - Set resistance_mechanism_class if the paper indicates the closest known
    homolog family (e.g. "aminoglycoside acetyltransferase")
  - Genes that were computationally predicted but NOT experimentally validated
    in this paper → do NOT extract

================================================================================
SECTION 8 - OUTPUT STRUCTURE
================================================================================
STRUCTURAL RULE: Gene entries NEVER carry a "mutations_found" field.

{
  "pmid": "<pmid string>",
  "relevant": true | false,
  "paper_title": "<title>",
  "publication_year": "<year>",
  "paper_type": "single_gene" | "multi_gene_resistome" | "review" |
                "computational_with_validation",
  "genes": {
    "<gene_name>": { ...GENE_ENTRY_SCHEMA fields only - NO mutations_found here... }
  },
  "mutations": {
    "<target_gene>": {
      "encodes": "what type of product this gene encodes",
      "mutations_found": [ { ...MUTATION_ENTRY_SCHEMA fields... } ],
      "wild_type_susceptible": "yes" | "no" | null,
      "notes": null | "<string>"
    }
  },
  "sequence_accessions": [ "<accession>", ... ],
  "key_findings": "<1-2 sentence summary of paper's main AMR findings>",
  "methodology": "<1 sentence on experimental methods used>",
  "geographic_location": [ "<region>", ... ],
  "sample_size": "<string>"
}

FIELD NOTES:
organisms_tested_in should almost never be empty. Look for:
  - The bacterial species the gene was cloned or isolated from (check
    the paper title, abstract, and Methods - it is almost always there)
  - The expression host used for MIC testing (commonly E. coli)
  - Any species named in susceptibility or MIC testing
  Use standard binomial nomenclature with serovar/strain where given
  (e.g. "Salmonella enterica serovar Typhimurium DT104", not just
  "Salmonella"). If the gene is from organism A and tested in organism B,
  list both. Only leave empty if the paper truly names no organism at all.

If the paper is not AMR-relevant at all, return:
{"pmid": "<pmid>", "relevant": false, "genes": {}, "mutations": {}}

================================================================================
SECTION 9 - SELF-CHECK (run before output)
================================================================================
Step 0 - GATE CHECK
 DIRECTION CHECK (run before everything else):
Ask: "Does this gene/mutation REDUCE susceptibility of a microorganism to
an antimicrobial agent?"
  YES → proceed to other checks
  NO, or the effect is INCREASED susceptibility / drug toxicity /
  sensitization → remove from output entirely

SUBJECT CHECK:
Ask: "Is the resistance or susceptibility phenotype measured in a
microorganism (bacterium, fungus, parasite)?"
  YES → proceed
  NO (phenotype is in human cells, animal tissue, mitochondria of
  eukaryotic cells, hair cells, etc.) → remove from output entirely

  Step 0a - GATE CHECK each entry in "genes":
  (a) Does the gene name belong to a recognized AMR family OR is it a validated
      gene in the papers?
  (b) Was the gene experimentally validated in THIS paper?
  (c) Does the wild-type confer resistance without mutation?
  If any answer is NO → remove from "genes" (move to mutations if appropriate).

Step 1 - Results-only check: every gene/mutation must appear in Results, Methods
(as cloned/assayed in this study), Tables, or Abstract findings.

Step 2 - Drug-class check: each confers_resistance_to entry must be consistent
with the gene family.

Step 3 - Hallucination check: any codon_change, nucleotide_change, or position
not literally written in the paper → set to null.

Step 4 - Table coverage: did you scan every table? Recount.

================================================================================
PAPER TEXT
================================================================================
```

{paper_text}

**S5.9 Auditor prompt (Pass 2)**

Reproduced as executed. {entity_type}, {entity_name}, {entity_json} and {paper_text} are runtime placeholders. One audit call was issued per extracted entity.

```
You are a strict AMR data auditor. You are given:
  1. The full text of a paper
  2. A single extracted entity (gene OR mutation target) as a JSON object

YOUR JOB: For EACH field in the JSON object, point to the EXACT sentence in the
paper that supports the value. If you cannot find explicit support, SET the field
TO NULL (do not omit it).

OUTPUT: a single JSON object containing the audited entity. Start with { immediately.
No prose, no markdown, no explanation. If the entity should be removed entirely
(no support in paper, or misclassified), return an empty object: {}.

RULES:
  - MISCLASSIFICATION CHECK (CRITICAL): Acquired AMR "genes" confer resistance in their wild-type form just by being present.
    * If ENTITY TYPE is "Gene", you MUST return {} to completely remove it if the paper implies ANY of the following:
      a) It is an "intrinsic", "baseline", or "endogenous" resistance determinant.
      b) Disrupting, knocking out, or mutating the gene causes SUSCEPTIBILITY, "supersusceptibility", or an "MIC reduction".
      c) Mutations, overexpression, or loss-of-function are required to gain resistance.
      d) A gene where the text specifies that "mutations", "amino acid changes", or "truncations" are what confer resistance.
      e) It is a core structural/biosynthesis gene (e.g., LPS, cell wall, glycosyl transferase) or a general regulatory gene (e.g., response regulator, two-component system).
      f) The paper's only experimental validation of this gene involves
         virulence, pathogenesis, or host-interaction phenotypes (macrophage
         survival, apoptosis, invasion, LD50, animal models) with no
         antimicrobial susceptibility data directly attributed to this gene.
         A gene validated only for virulence is NOT an AMR gene.

  - Find the specific section in the papers to check each field validity. SET fields to NULL when unsupported. Do not delete them.
  - Set codon_change, nucleotide_change, nucleotide_position to NULL UNLESS the paper
    explicitly writes them (e.g. "AGG→TAT", "G249T", "position 249"). Verbatim
    only - do not infer from amino-acid change.
  - For drug classes: if the papers does not state specifically that this gene family confers resistance to a given
    antibiotic, remove that antibiotic. Do not rely on inference from family name alone.
  - organisms_tested_in / organisms_observed_in: This field should almost
    never be empty. Actively search the paper for:
      * The species the gene was cloned or isolated FROM (e.g. "from
        Salmonella typhimurium DT104")
      * The host used for functional testing (e.g. E. coli transformants)
      * Any species named in MIC testing, disk diffusion, or susceptibility
        assays
    If the paper studies a gene from organism A expressed in organism B,
    list both. Use standard binomial nomenclature with strain/serovar where
    given (e.g. "Salmonella enterica serovar Typhimurium DT104"). Check the
    title, abstract, and Methods section - the organism is almost always
    named in at least one of these. Only leave empty if the paper truly
    does not name any organism.

  - For mutation targets: if the paper does not explicitly state that "mutations in X" or "inactivation of X" confers resistance
    to the specified drug, remove the drug from confers_resistance_to. Do not rely on inference from family name alone.

ENTITY TYPE: {entity_type}
ENTITY NAME: {entity_name}
ENTITY DATA:
{entity_json}

PAPER TEXT:
{paper_text}
```

AUDITED JSON OBJECT:

**Section S6. Harmonization and tiering**

This section documents how ResLit records were harmonized against curated reference data and assigned evidence tiers. Two pipelines are assembled independently. The first one is the ResLit extraction pipeline described in Sections S1-S5; and the second is a reference pipeline drawing on CARD, ResFinder, and the NCBI Reference Gene Catalog. The per-step processing chains for genes and mutations are given below, followed by the tier logic.

**S6.1 Reference-database assembly**

Curated reference records were assembled from three sources. The NCBI Reference Gene Catalog was obtained from the AMRFinderPlus database distribution) and filtered to AMR core-scope entries, with protein accessions fetched individually. CARD was downloaded from its distribution portal (aro_index, card.json, sequence FASTAs, SNP definitions, and PMID mappings), and entries scoped only to disinfecting agents and antiseptics were removed. ResFinder (version 2.6.0) was obtained as a versioned repository clone and translated to protein; point-mutation definitions were drawn from a PointFinder export.

Protein sequences from the three sources were pooled and clustered at 100% identity using the toolCD-hit. Records clustering together across sources were merged into a single entry carrying the union of their database memberships, so that a determinant curated by more than one database is represented once with each contributing source recorded. This produced a single reference gene table; reference mutation records were assembled separately from PointFinder, the NCBI point-mutation catalogue, and CARD SNP definitions, reshaped into a common schema and classified as nucleotide- or amino-acid-level changes.

**S6.2 ResLit gene post-processing chain**

The per-batch extractor output (Section S5) was converted to tabular form and processed through the following ordered chain. Row counts are approximate and specific to the current release.

| **Step** | **Operation** |
| --- | --- |
| 1 | Convert extractor JSON to per-batch CSV; canonicalise gene/allele names against the shared gene-family reference (allele-family map), fix missing bla prefixes and aminoglycoside prime-mark notation, and drop locus tags, hallucinated repeated-character strings, and non-bacterial gene names. |
| 2 | Concatenate the six batches into a single table. |
| 3 | Retain only rows whose evidence level indicates experimental characterisation. |
| 4 | Add a lexically normalised gene-name column (lowercase, prime/quote marks stripped) for matching. |
| 5 | Drop rows lacking a paper title; label every row with source “Reslit”. |
| 6 | Select into the final gene schema ( target schema: Database, Gene, Allele, Encodes, Mechanism, Resistance, Organism, accessions, Validation_method, PMID, and paper metadata). |
| 7 | Harmonise gene-name capitalisation against the reference catalogue (S6.4). |
| 8 | Filter to recognised bacterial organism and recognised antibiotic (S6.5), yielding the seed gene file. |

**S6.3 ResLit mutation post-processing chain**

Mutations were processed through a parallel chain. Unlike the gene chain, the mutation chain includes no cross-database name harmonisation step; reference comparison for mutations occurs only at database load (S6.6).

| **Step** | **Operation** |
| --- | --- |
| 1 | Convert extractor JSON to per-batch CSV. The mutation converter additionally special-cases non-coding RNA loci (16S/23S, rrs/rrl), expands compound mutations into one row each (e.g. A90V + D94G → two rows), and requires a recognised antibiotic unless the entry is a promoter mutation. A hallucination-detection regex was corrected during development from a form that matched almost any string to a backreference-based repeated-character check. |
| 2 | Concatenate the six batches. |
| 3 | Enrich rows with missing title/year: fetch by PMID where present, or resolve a PMID by searching PubMed on the row's sequence accession, then fetch metadata. |
| 4 | Retain only rows with a non-empty normalised gene and normalised change. |
| 5 | Drop changes containing ambiguous characters, reconcile rrs/rrl mislabels, and require a reported organism. |
| 6 | Select into the final mutation schema, expanding each pipe-separated organism into its own row (≈26,148 rows). |
| 7 | Fix gene-name capitalisation. |
| 8 | Filter to recognised bacterial organism and recognised antibiotic (S6.5), yielding the seed mutation file (26,094 rows). |

**S6.4 Name harmonisation (genes only)**

Harmonisation aligns ResLit gene names to the capitalisation used in the reference catalogue. The combined reference file is treated as the canonical naming authority: for each normalised gene name, its most frequent original-case spelling in the reference data is adopted, and ResLit Gene and Allele fields are rewritten to match wherever a normalised match exists. This step is purely cosmetic - no rows are added, dropped, or content-transformed, and applies to the gene catalogue only; the mutation catalogue receives capitalisation fixes but no cross-database name alignment. A companion capitalisation pass corrects names against a curated bacterial gene-name list.

**S6.5 Organism and antibiotic filtering**

The ResLit gene and mutation catalogues were each filtered to entries carrying a recognised bacterial organism and a recognised antibiotic, with both fields standardised during the process. Organism names were checked against a controlled species list (with a small set of higher-taxon and common names); unrecognised names were cleared from the cell. Antibiotic names were checked against controlled antibiotic-name and abbreviation lists, with plural, hyphen, and slash normalisation and de-duplication. Rows left with no recognised antibiotic were dropped, with one exception: promoter mutations, identified by negative nucleotide coordinates, were exempt from the antibiotic requirement. Mutation entries additionally underwent a nucleotide-versus-protein position consistency check, in which the nucleotide position was required to be approximately three times the protein position or the nucleotide change was cleared.

**S6.6 Database load and evidence tiering**

The two ResLit seed files and the two reference seed files were loaded into a relational database, together with a papers table derived by de-duplicating PMIDs across all four. Reference-database source labels were normalised to a common set (Reference Gene Catalog, CARD, ResFinder) so that each ResLit determinant could be compared against its presence in each reference resource. On the basis of this comparison, every gene and mutation was assigned one of four evidence tiers defined by the intersection of two independent lines of evidence - presence in a curated reference database, and recovery by ResLit from the primary literature.

| **Tier** | **Definition** |
| --- | --- |
| Confirmed | Corroborated by more than one independent line of evidence: either present in two or more reference databases, or present in one reference database and independently recovered by ResLit. |
| Established | Curated in one reference database but not recovered by ResLit, whether because the source publication lies outside the retrieved corpus or because extraction did not identify the association. |
| Supported | Absent from all three reference databases, but reported in three or more independent papers in the ResLit corpus. |
| Candidate | Absent from all three reference databases and reported in fewer than three papers. |

Curator accounts, comments, and curation history are stored separately and are not sourced from the seed files; re-seeding the catalogues leaves curatorial data untouched.
